# Dynamics of Contaminant Microbes in Bioethanol Production from Sugarcane

**DOI:** 10.64898/2026.02.04.703784

**Authors:** Ana Laura Ramos Romano, Natalia Coutouné, Artur Rego-Costa, Michael M. Desai, Marcelo F. Carazzolle, Andreas K. Gombert

## Abstract

The dynamics and impact of microbial contaminants in industrial sugarcane bioethanol production in Brazil were investigated through a two-year metagenomic study across two biorefineries. Shotgun metagenomic sequencing revealed that temporal shifts in the contaminant microbiome dynamics within production seasons were more pronounced than inter-annual or inter-mill variations. While *Saccharomyces* spp. dominated, bacterial communities, primarily within the Firmicutes phylum and dominated by the genera *Lactobacillus*, *Limosilactobacillus*, and *Bacillus*, exhibited dynamic changes. Correlation analyses with industrial process parameters revealed a complex interplay: lower *Lactobacillus* levels in one mill were associated with increased ethanol yield, whereas higher levels in another mill correlated with reduced yeast viability and increased flocculation. The presence of *Limosilactobacillus* was linked to decreased yeast viability, whereas *Bacillus* showed potential for inhibiting both *Lactobacillus* and *Limosilactobacillus*. These findings highlight the nuanced and species-specific impacts of bacterial contaminants on bioethanol production, underscoring the need for strain-level functional studies and targeted interventions to optimize fermentation efficiency and stability in industrial settings.

## INTRODUCTION

The global push for sustainable energy positions bioethanol as a key renewable fuel. Brazil is a leader in this market, ranking as the world’s second-largest producer and top exporter (1)(2). Its sugarcane-based process is vastly more land– and energy-efficient than the corn-based method used by the largest producer, the United States (3)(4). Brazilian sugarcane bioethanol boasts an energy balance of 10.2:1 (renewable output to fossil input), while US corn bioethanol is significantly lower at roughly 1.4:1 (5). This production relies on the fermentation of sugarcane feedstocks by *Saccharomyces cerevisiae* in non-aseptic conditions, making it highly susceptible to microbial contamination and related process disruptions. These contaminants, primarily bacteria, may compete directly with *S. cerevisiae* for sugars, diverting resources that would otherwise be transformed into ethanol, reducing the final achievable ethanol yield. In an industry where profit margins are exceptionally tight, even minor gains in process efficiency can have a substantial economic impact.

The sugarcane bioethanol industry employs multiple strategies to limit microbial contamination in the fermentation environment. This includes prophylactic sulfuric acid treatment of recycled yeast cells (pH < 2.0 for 1 to 1.5 h) to suppress bacterial growth (7), maintaining a high yeast inoculum fraction, and performing routine equipment cleaning to reduce biofilm formation. Although antibiotics such as penicillin, virginiamycin, streptomycin, tetracycline, and monensin have historically been used, rising concerns over increased antibiotic resistance and potential contamination of animal feed byproducts (8) have prompted the search for alternative antimicrobial approaches. Bacterial contamination levels can reach 10⁷–10⁸ cells/mL, leading to economic losses of up to 20,000 liters of bioethanol per day in a medium-sized biorefinery (9), primarily due to competition for nutrients and the production of inhibitory organic acids (10, 11). Moreover, species like *Limosilactobacillus fermentum* can cause yeast flocculation (12), disrupting yeast–substrate interactions, slowing fermentation kinetics, and reducing ethanol yield and productivity, while also impairing downstream processes such as yeast separation before distillation (13).

Despite the economic and operational significance of bacterial contamination in bioethanol production, our understanding of the dynamic shifts within these bacterial communities over time remains limited. Culture-dependent methods have identified key genera like *Lactobacillus, Bacillus, Staphylococcus,* and *Leuconostoc* (14–17), and more recent studies using 16S rRNA gene sequencing consistently highlight *Lactobacillus* as the dominant contaminant (10, 11, 18–21). However, these reports provided only a low-resolution and rather static snapshot of microbial community composition, because they relied on a small number of samples, making it difficult to establish correlations between microbial composition dynamics and process performance (22).

Shotgun metagenomic sequencing offers a high-resolution alternative to observe microbial communities. Lino et al. (2024) applied this strategy to samples from two sugarcane biorefineries, revealing a more diverse microbial composition in the fermentation environment than previously described, and proposing new hypotheses about the impact of some of the bacterial members on the industrial process (23). However, such datasets are still scarce, especially when sampling across different production cycles and biorefineries. Further lack of temporal resolution prevents evaluation of the population dynamics of these microbial communities.

To address this knowledge gap, in this study we use whole-metagenome sequencing to characterize the microbiome present in the industrial fermentation of sugarcane feedstocks. We evaluated the temporal changes in this microbiome across two entire production years in two distinct biorefineries, expanding upon the initial findings of Rego-Costa et al. (24) by specifically analyzing DNA sequences not attributable to *S. cerevisiae*. This approach aims at providing a deeper understanding of the bacterial dynamics within industrial alcoholic fermentations in Brazilian sugarcane biorefineries, ultimately contributing to the development of more effective strategies for contamination control and process improvement.

## MATERIALS AND METHODS

### Sample Collection

Industrial fermentation samples were collected approximately weekly over two consecutive sugarcane production seasons (2018 and 2019), spanning from April to November of each year. Samples were obtained from two independent sugarcane mills, designated as UIR and UCP (Fig. 1A, corresponding to Site A and Site B, respectively, in (24)), both situated in the region of Piracicaba, São Paulo State, Brazil. All samples were taken from the fermented broth, or *wine*, in the industrial jargon, either directly from a fermentor or from the holding tank used as an intermediary storage vessel downstream of the fermentor and upstream of the continuous centrifuges. For each sampling event, 15 mL Falcon tubes prefilled with 3 mL of sterile glycerol were filled with 12 mL of the sample. Immediately after collection, samples were thoroughly homogenized by vortexing and then subjected to a two-stage freezing protocol: initial storage at –20 °C for 1 to 3 months, followed by long-term storage at – 80 °C. Samples were later shipped in dry ice to Harvard University, where they were kept at –80 °C until further processing. The collection and handling of these samples were conducted in accordance with Brazilian federal regulations, and the samples are duly registered in the SisGen database under the accession numbers R40E57A, RB42674, R193AED, RAD5521, and AF14971.

**Figure 1.**
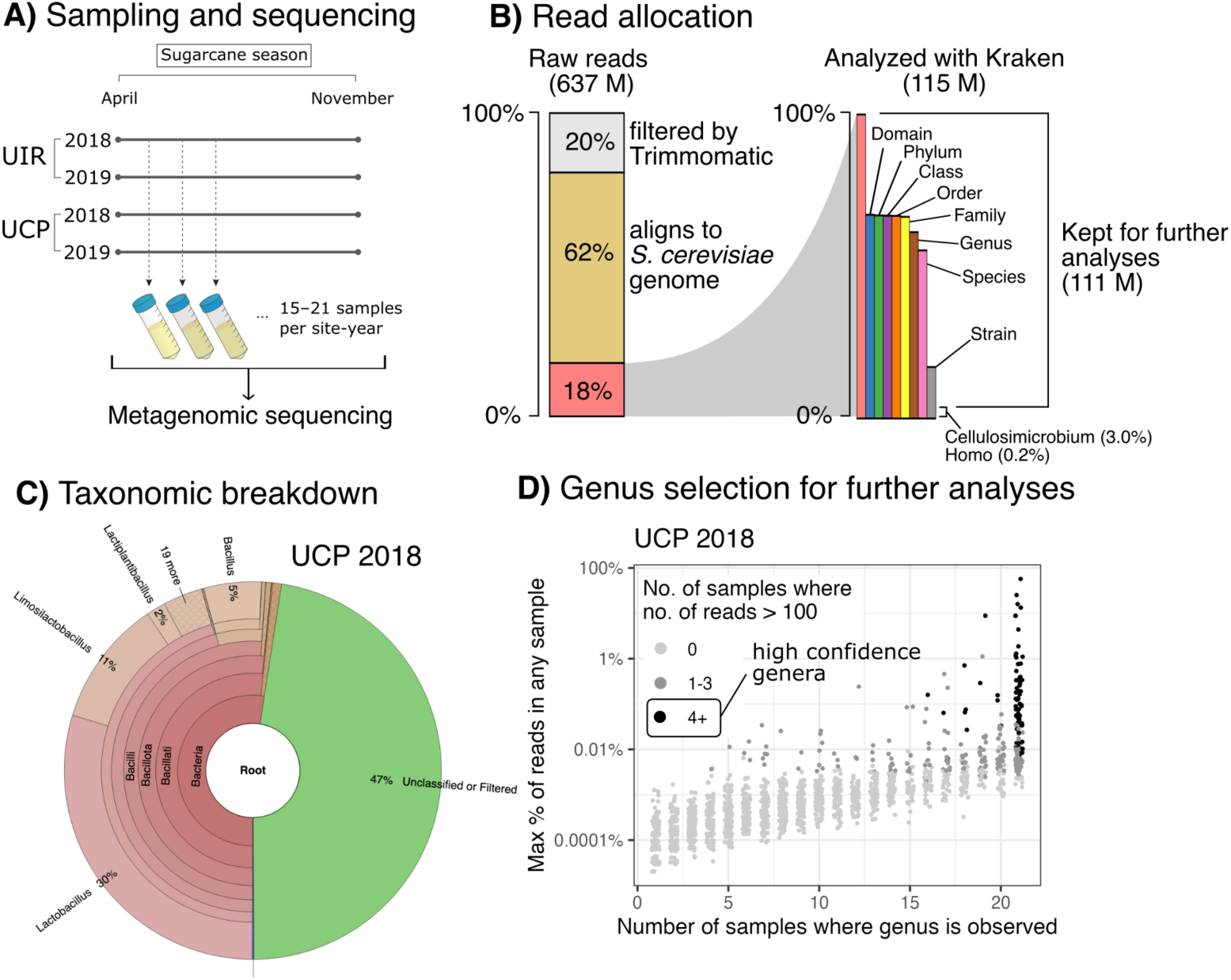
Metagenomic pipeline and selection of high-confidence genera for bioethanol fermentation contaminants. (A) Schematic of the time-series sampling strategy across two biorefineries (UIR and UCP) during the 2018 and 2019 sugarcane seasons, leading to shotgun metagenomic sequencing. (B) Read processing and allocation pipeline. Raw reads were filtered by quality (20%), and budding yeast-derived sequences were removed by alignment to the s288c reference genome (62%). The remaining high-quality reads (18%) were classified using Kraken2, with the fraction of classified reads decreasing at more granular taxonomic levels. See a breakdown of this classification in each site-year and across temporal samples in Supplementary Figure 1. (C) Illustrative taxonomic breakdown of classified reads in UCP 2018, showing dominance of bacterial taxa, notably Lactobacillus (30%) and Limosilactobacillus (11%). This data is available as an interactive plot in Supplementary Material 2. (D) Genus selection methodology (in UCP 2018) based on peak observed abundance (maximum fraction of reads in any sample) and prevalence (number of samples where genus is observed). High-confidence genera (black points) were retained for further analysis based on the criterion of >4 samples containing >100 reads. See the same figure for the other three site-years in Supplementary Figure 2.

### DNA Extraction and Sequencing

A total of 70 samples collected from both mills and both production years were selected for shotgun metagenomic sequencing. Information on sample collection dates can be found in Rego-Costa et al. (2023) (24). Total genomic DNA was extracted from each sample using a column-based protocol previously described by Nguyen Ba et al. (2019) (25), which was assessed visually under a microscope to ensure efficient lysis of all cells in the samples. Sequencing libraries were prepared using a transposase-based kit (Illumina, San Diego, CA, USA) following the reduced-volume protocol described by Baym et al. (2015) (26). Library quality and size distribution were assessed using an Agilent TapeStation 4200 (Agilent Technologies, Santa Clara, CA, USA), and libraries were quantified using Thermo Fisher’s QuBit dsDNA HS assay before sequencing. Sequencing was performed using two runs on an Illumina NextSeq 500 platform and one run on an Illumina MiSeq platform, generating an average of 9.1 million 300 bp paired-end reads per sample.

### Read Preprocessing

Raw paired-end reads were trimmed using Trimmomatic version 0.39 (27), removing the reads shorter than 36 base pairs and trimming of low-quality bases from the 3’ end of reads using a sliding window of 4 bases, with a minimum average quality score of 23, which removed ∼20% of all the reads (Fig. 1B). Adapter sequences were removed using NGmerge version 0.2 (28) with the “-a” flag, employing stringent parameters to ensure complete adapter removal and accurate merging of overlapping paired-end reads, where applicable.

### Taxonomic Profiling

Taxonomic classification of the processed reads was performed using Kraken 2 (v. 2.0.8-beta) (29), a *k*-mer-based classifier known for its speed and accuracy. Initially, all reads were classified at a high taxonomic level (kingdom) against the core_nt database available from the Kraken 2 repository (https://benlangmead.github.io/aws-indexes/k2) to delineate reads belonging to *Saccharomyces*, other fungi, bacteria, viruses, and archaea. Due to the overwhelming abundance of *Saccharomyces cerevisiae* reads, which could obscure the detection of less abundant microbial taxa, a targeted approach was employed to remove these *Saccharomyces*-derived sequences. All processed reads were aligned against the reference genome of *Saccharomyces cerevisiae* S288C using the Burrows-Wheeler Aligner (BWA) version 0.7.15-r1140 (30). Reads that failed to align to the *S. cerevisiae* genome were retained for subsequent taxonomic classification of the remaining microbial community (Fig. 1B; Supp. Figure 1). These unmapped reads were then classified using Kraken 2 against its bacterial, fungal, viral, and archaeal databases. The raw output from Kraken 2 is included in Supplementary Material 1. The breakdown of the classification across all four site-years was visualized using Krona (Fig. 1C displays data for UCP 2018) (31). An interactive chart can be found in Supplementary Material 2.

Kraken 2 classifies reads down to the smallest taxonomic level that it can deal with statistical confidence. In our dataset, 67% of the reads could be classified to the coarsest taxonomic level, Domain, with 61% being classified down to Genus, followed by a dropoff to 55% at the level of Species (Fig. 1B). To balance classification depth and amount of classified reads, we retained all classifications down to the Genus level in downstream analyses. Given the inherent uncertainty in read classification, we employed a filter on classified genera to each site-year separately, only keeping high-confidence genera that are seen to have more than 100 reads in 4 or more samples in that site-year (Fig. 1D). We further removed any reads classified down to the genus *Homo*, which likely stem from contamination during any step of the process, as well as those classified down to *Cellulosimicrobium*, which are likely to stem from the enzymatic treatment used for yeast cell lysis in the genomic DNA preparation step. In all further analyses, we refer to the ratio between the number of reads classified down to a kept genus and the total number of reads fed to Kraken 2 (*Homo* and *Cellulosimicrobium* excluded) as either *Relative abundance* or *Fraction of Reads*.

### Microbial community analysis

Microbial community structure was explored using multiple complementary approaches. Principal component analysis (PCA) was carried out over the relative abundance data for both years and sites together using the factoextra package (32) in R (v.4.3.1). This provided us with a way to visualize global patterns of variation in microbial composition and assess potential clustering by industrial site and production year.

The alpha diversity of bacterial communities was assessed using the Shannon diversity index (H’). Shannon’s index was then computed as:

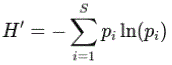

where *p_i_* represents the relative abundance of genus i among S genera detected in the sample (33). This index accounts for both richness and evenness of the microbial community, with higher values indicating greater diversity.

To investigate potential interactions within the microbial community, we inferred co-occurrence and co-exclusion networks from genus-level abundance data. Networks were constructed using the SpiecEasi package (34) with the Meinshausen–Bühlmann method and 99 regularization parameters, with model stability evaluated by pulsar subsampling (50 replicates). The resulting adjacency matrix of partial correlations was represented as an undirected weighted graph using igraph (35). Edges above the 80^th^ percentile of absolute weight were retained and classified as positive (co-occurrence) or negative (co-exclusion) interactions. Final networks were visualized with ggraph (36) using a Fruchterman–Reingold layout, with node size scaled to degree and edge width proportional to correlation strength.

### Industrial process metrics and regression analysis

Industrial process data were obtained from both sites and encompassed a range of fermentation and yeast performance variables (Supp. Table 1). These included yeast-related metrics (viability, budding, flocculation, and bacterial counts) and physicochemical parameters of the fermentation process (sugar concentration in must and wine, pH, fermentation yield, and volume of ethanol produced). The frequency and type of measurements differed between the two sugarcane mills: UIR performed detailed yeast viability and contamination monitoring, while UCP reported physicochemical data and weekly yield records. A detailed description of individual parameters can be found in Supplementary Table 1.

To test for associations between microbial community variation and industrial performance, we implemented linear regression models linking the abundance of the two main genera to occupy the process: *Lactobacillus* and *Limosilactobacillus* to each industrial variable. For each site (UIR and UCP), separate models were fitted with the general form:

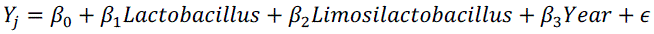

where Yj represents each industrial metric j (e.g., yeast viability, pH, ethanol yield), and Year was included as a discrete covariate to account for interannual effects. Models were fitted using the lm() function in R, and p-values for regression coefficients were computed from t-tests to assess the statistical significance of each association.

### Data and code availability

The raw sequencing data have been deposited in the NCBI Sequence Read Archive (SRA) under the BioProject accession number PRJNA865262. All code developed and used for metagenomic processing and statistical analyses is publicly available in the GitHub repository: https://github.com/alrsilva5577/Contaminants_ethanol.

## RESULTS

This study involved the collection of whole-population microbiological samples during the 2018 and 2019 fermentation seasons from two independent sugarcane biorefineries, referred to as UIR and UCP (Fig. 1A). These mills are located approximately 18 km apart in São Paulo state, Brazil. Operational differences — such as the yeast starter strains used at the beginning of the season and distinct contamination control strategies, among others — can influence the microbiological community structure of the fermentation process. Previous work by Rego-Costa et al. (24) described the dynamics of the yeast *S. cerevisiae* populations in these same samples, showing large population shifts, including the invasion of the process by new yeast strains likely originating from the environment. Here, we describe and analyze the temporal dynamics, microbial community structure, and correlation with industrial process metrics of the total microbial DNA excluding *S. cerevisiae*.

Shotgun metagenomic sequencing of the 70 collected samples yielded a total of 639 million 300-bp paired-end reads, with an average of 9.1 million reads per sample (Fig. 1B). 20% of the reads were first removed due to low quality scores based on the established Trimmomatic filtering criteria. Approximately 62% of all reads aligned to the *Saccharomyces cerevisiae* S288C reference genome. The remaining 18% of high-quality, non-*Saccharomyces* reads were subjected to taxonomic classification with Kraken 2 in order to characterize the microbial contaminants present in the fermentation environment, including archaea, viruses, bacteria, and other fungi.

The fraction of reads classified to each taxonomic level was consistent across all site-years and time points (Fig. 1B; Supp. Fig. 1). Given the probabilistic nature of taxonomic assignment and the sharp drop-off in fraction of classified reads when comparing genus to species, we proceeded with genus-level classification data. That is, reads classified down to the genus or lower taxonomic level retain their genus classification, and reads classified down to a taxonomic level above genus remain labeled as Unclassified at the genus level. We further reassess the genus-level classification. We start by removing the reads identified as likely contaminants: *Homo*, likely from human handling in both the industry and in the laboratory, and *Cellulosimicrobium*, likely related to the *Cellulosimicrobium*-derived zymolyase enzymatic treatment used in genomic DNA preparation. We then leveraged the longitudinal sampling to filter for high-confidence genera, retaining only those genera present with >100 reads in at least four samples within a given site-year (Fig. 1D; Supp. Fig. 2). This high-confidence set accounted for 94% of all reads classified to the genus level; reads from filtered-out genera were relabeled as Unclassified. We use this data for all further analyses, using read abundance (the fraction of all high-quality non-*Saccharomyces* reads classified to a given genus) as a proxy for population abundance that we do not attempt to correct for genome size.

**Figure 2.**
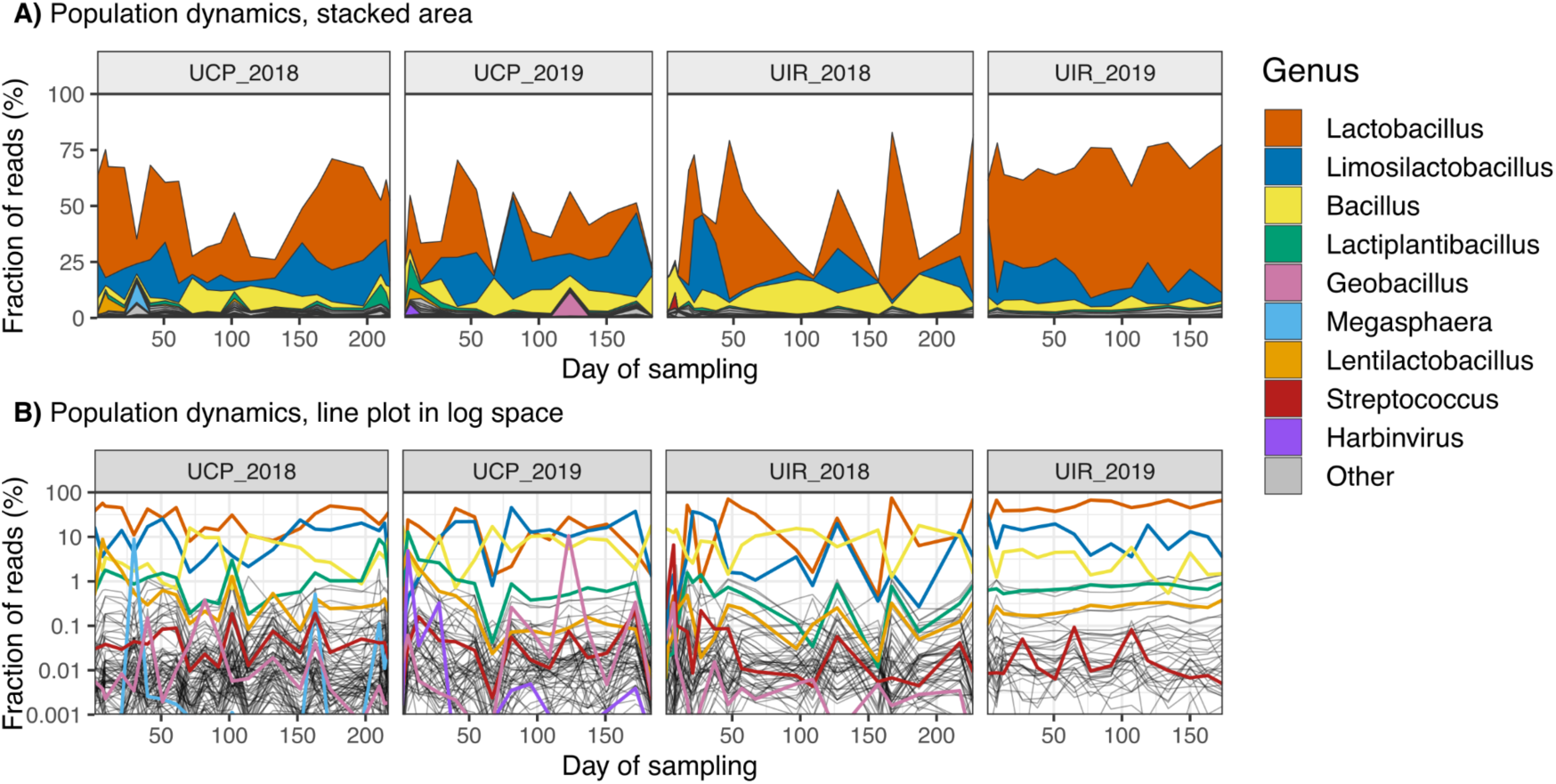
Temporal dynamics of the microbial (non-*Saccharomyces*) community in industrial bioethanol production. Relative abundance of identified bacterial genera tracked over the sampling period for all four site-years. (A) Stacked area plots showing community composition, dominated by *Lactobacillus* and *Limosilactobacillus*, followed by *Bacillus*. The top nine most abundant genera are distinctly colored, while the remainder are shown in grey. (B) Line plots of the same genus abundance data, displayed on a log-scale (0.001% to 100% fraction of reads). This visualization highlights the temporal fluctuations and persistence of low-abundance lineages across all four-time series.

The final filtered dataset comprised 98 unique bacterial genera and 5 bacteriophages (Supp. Fig. 3). Consistent with literature on the bioethanol industry (37), the communities were consistently dominated by the genera *Lactobacillus*, *Limosilactobacillus*, and *Bacillus*, which were the most prevalent genera across all datasets (Fig. 2; Supp. Fig. 3). The overall community structure was heavy-tailed and showed site-specific differences (Supp. Fig. 4). UCP 2018, for example, exhibited the highest richness with 81 different genera, 24 of which are unique to this site-year, while UIR 2019 was the least rich, with 36 total genera, and a single unique genus (Supp. Fig. 4A; Supp. Fig. 5). Beyond the dominant taxa, we identified a persistent community of low-abundance microbes, with 30 genera shared across all four site-years, underscoring their stable role in this fermentation environment (Supp. Fig. 4A).

**Figure 3.**
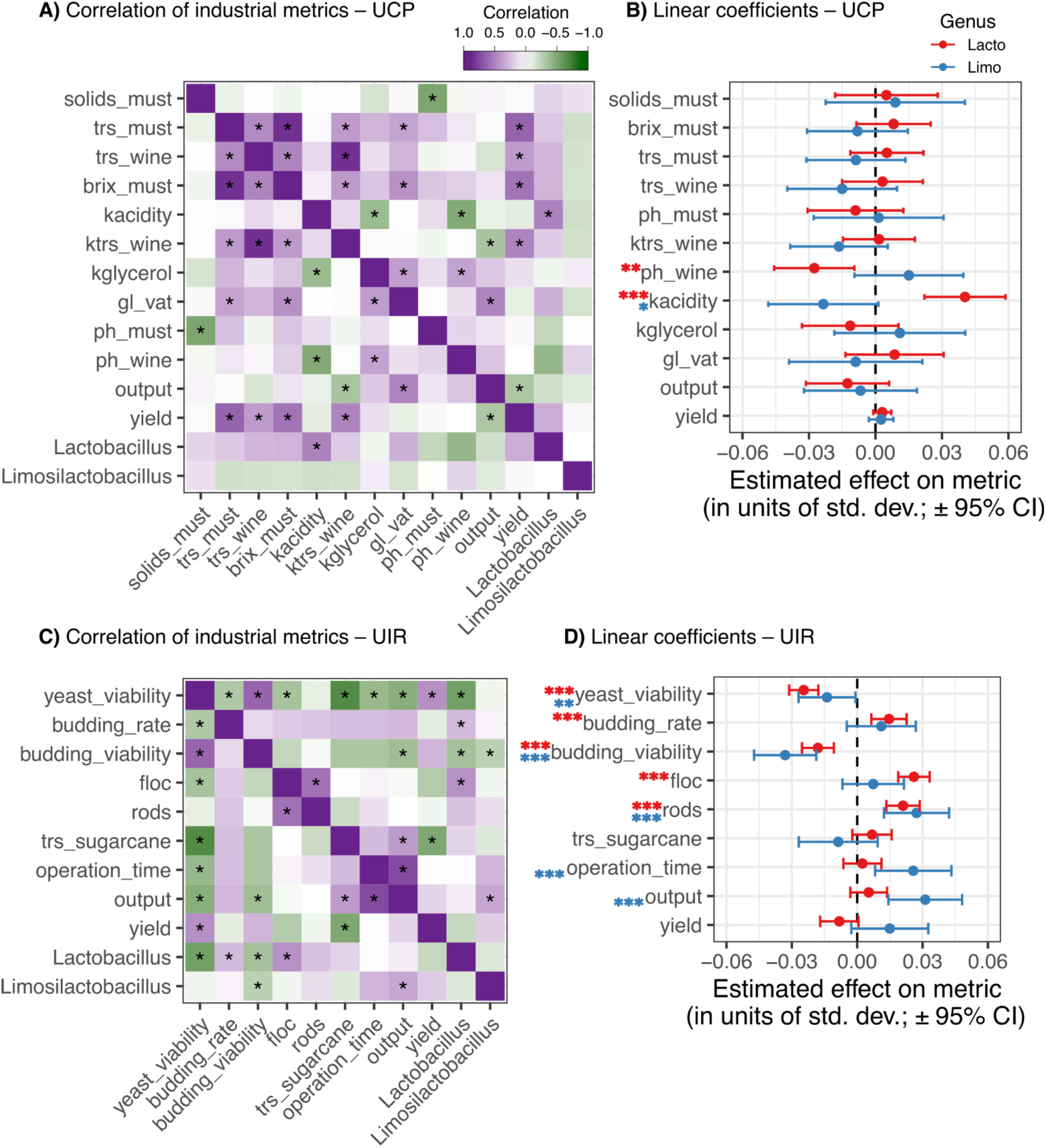
Statistical association of dominant bacterial genera with industrial process metrics. (A, C) Pearson’s correlation matrices for UCP and UIR, respectively, showing correlations between genus abundance and metrics of the industrial process, kindly shared with us by the biorefineries. See Supplementary Tables 1, 2, and 3 for more information on these metrics and for the fit statistics of the regression models. Asterisk indicates a significant correlation under a t-test of the correlation coefficients: *: *p* < 0.1, **: *p* < 0.05, ***: *p* < 0.05/*no*. *of tests* as a Bonferroni-corrected significance threshold, where the number of tests is taken to be 12 for UCP and 9 for UIR. (B, D) Estimated linear coefficients and 95% confidence intervals for the effect of *Lactobacillus* and *Limosilactobacillus* abundance on industrial metrics at UCP and UIR, respectively, taking year as a covariate. Observed values and predictions from linear models are shown in Supplementary Figures 9 and 10.

Analysis of the temporal dynamics revealed the highly unstable nature of this microbial ecosystem (Fig. 2). The dominant genera, particularly *Lactobacillus* and *Limosilactobacillus*, exhibited large boom and crash cycles, with relative abundances fluctuating dramatically between successive time points (Fig. 2A). An exception was observed in the UIR 2019 time series, which showed a qualitatively different dynamic. Here, the community composition remained relatively stable throughout the season, a feature that also corresponded to the lowest overall genus richness and Shannon diversity of all four datasets (Supp. Fig. 5). While *Lactobacillus* and *Limosilactobacillus* abundance fluctuations did not show a clear relationship, their cycles often coincide with fluctuations in multiple low-abundance genera, a pattern suggestive of system-wide perturbations and/or microbial interactions (Fig. 5B).

We next sought to determine if the observed microbial dynamics were associated with industrial process metrics (Supp. Table 1; Supp. Fig. 6, 7). The specific metrics collected varied between the UIR and UCP facilities. As expected, many of these metrics correlate with one another (Fig. 3A,C), including trivial associations (e.g., between total reducing sugar, or TRS, and the Brix index in UCP) and non-trivial ones (e.g., yeast viability and sugarcane TRS in UIR).

To assess the impact of the dominant bacteria in the process, we focused on *Lactobacillus* and *Limosilactobacillus*. These two genera are the most abundant and, critically, their abundances do not correlate, as assessed by Pearson’s correlation or by PCA (Fig. 3A,C; Supp. Fig. 8A), allowing their effects to be evaluated independently. We fit individual linear regression models for each industrial metric, using the abundances of both genera as explanatory variables and including the production year (2018/2019) as a covariate to account for annual variation.

The models revealed significant associations between bacterial abundance and process metrics at both facilities (Fig. 3B,D; Supp. Table 2,3). At UCP, *Lactobacillus* abundance showed a strong, statistically significant association with lower wine pH (i.e., higher acidity), a finding supported by both direct pH tests and titration methods (Fig. 3B). Conversely, *Limosilactobacillus* abundance was associated with a less acidic environment (a positive coefficient for pH). No other significant relationships were found at UCP between these two genera and other metrics, total ethanol output or production yield being relevant ones.

At UIR, the associations revealed a more complex interplay. Both *Lactobacillus* and *Limosilactobacillus* were significantly associated with decreased yeast cell and bud viability (Fig. 3D). *Lactobacillus* abundance was also correlated with a higher budding rate and increased flocculation, the latter referring to a state of cell aggregation associated with reduced fermentation efficiency. Most notably, *Limosilactobacillus* showed a significant positive association with key production metrics: Operation Time (the fraction of time the pipeline is operational) and Final Output (total ethanol volume). This suggests that while both dominant genera may negatively impact yeast health, *Limosilactobacillus* is linked to more favorable, large-scale industrial outcomes.

The co-occurrence network exhibited multiple interconnected taxa forming small clusters, mostly in pairs. Only a few nodes showed a higher degree, acting as local hubs, suggesting taxa with broader association profiles within the community (Supplementary Figure 11). Several genera that emerged from the recent reclassification of the former genus *Lactobacillus* (38) (e.g., *Companilactobacillus, Latilactobacillus, Liquorilactobacillus, Loigolactobacillus,* and *Paucilactobacillus*) formed a subnetwork of positive associations. These genera share a common evolutionary origin and are adapted to carbohydrate-rich, fermentative environments, suggesting that their co-occurrence reflects shared ecological preferences rather than direct biological interactions. Most of the other co-expression clusters (such as *Corynebacterium, Polynucleobacter*, and *Rhizobium*) represent a functional module of plant-associated microorganisms, linked by an ecological niche in the rhizosphere and phyllosphere, that persist after sugarcane processing.

While all identified viral taxa were characterized as bacteriophages of the *Lactobacillus* genus, two of them exhibited positive correlations with non-host bacterial genera (*Colunavirus* and *Watanabevirus*), and only the genus *Harbinvirus* was positively associated with the genus *Lactiplantibacillus* within the co-expression networks. Notably, no co-exclusionary patterns were observed between these phages and any of the newly reclassified *Lactobacillus* genera. This lack of negative correlation could suggest that viral activity in this ecosystem may not be driven by acute lytic predation of the dominant lactic acid bacteria, but rather by stable lysogenic associations as previously described for lactobacilli phages (40,41,42,43).

This pattern of small sub-networks, rather than a single highly connected core, may suggest a wide diversity of origins for the bacteria found in the fermentation tank. Many of the genera described are commensals or parasites of insects and plants such as *Apilactobacillus, Bombilactobacillus, Blattabacterium, Candidatum Nardonella, Spiroplasma,* and *Xanthomonas* (38, 44, 45, 46, 47, 52); others are organisms associated with the rhizosphere or freshwater such as *Bacillus, Pantoea, Mycobacterium, Flavobacterium* and *Chlamydia* (48, 49, 50, 51, 52), and several other genera with a high number of human or animals pathogens, like *Treponema, Haemophilus*, and *Yersinia* (53, 54, 55).

Interestingly, the co-exclusion subnetwork between *Bacillus* and *Lactobacillus* indicates a strong antagonism. As lactic acid producers, *Lactobacillus* species acidify the fermentation medium, creating unfavorable conditions that could inhibit *Bacillus’* growth and induce sporulation, as *Bacillus* prefers neutral to alkaline environments (48). This negative association could be further driven by the production of bacteriocins by *Lactobacillus*, a widely described phenomenon (56, 57, 58, 59, 60). Furthermore, the antimicrobial repertoire of *Bacillus* spp. spans numerous chemical classes, ranging from small non-ribosomal peptides to complex polyketides and lipopeptides, capable of producing quorum-quenching and causing cell lysis in other bacteria (61).

## DISCUSSION

Our study employed metagenomic sequencing to investigate the non-*Saccharomyces* fraction of the microbial communities in the economically-relevant fermentation process of bioethanol production from sugarcane. By sampling these communities with high temporal resolution across biorefineries and over different fermentation seasons, we were able to assess the consistent and variable aspects of the microbial composition and dynamics of this industrial process with higher confidence than has been possible so far.

We have found *Lactobacillus* and *Limosilactobacillus* to be the most prevalent contaminants of this industrial alcoholic fermentation process, which agrees with previous studies (22). While both genera produce lactic acid, they differ in their metabolic impact on the process. *Lactobacillus* is strictly homofermentative (38), producing virtually exclusively lactic acid as the product of glucose fermentation, while *Limosilactobacillus* is heterofermentative, producing ethanol and acetic acid, in addition to lactic acid. In UCP we have found that *Lactobacillus* abundance is associated with a more acidic fermentation environment, while the opposite is true for *Limosilactobacillus*. This contrasts with the observations of Lino et al. (2024) (37), who found *Limosilactobacillus* to be associated with higher acidity. Lino et al. (2024) sampled the process over the course of a fermentation cycle, while we sampled mostly the fermented broth at the end of a cycle over the course of whole fermentation seasons. Differences in temporal resolution may account for these contrasting observations. We also note that both studies are limited by the use of correlation-based analyses, which hampers our ability to make causal claims.

Biorefineries in our study differed in their antimicrobial strategy during the study period. In the 2018 production year, UIR initiated operations with a low dose of monensin, subsequently transitioning at some unknown timepoint to the natural antimicrobial hops and the chemical chlorine dioxide for the remainder of the year. In 2019, UIR switched to using exclusively hops and chlorine dioxide, with no antibiotic application. UCP utilized a combination of three antibiotics – monensin, virginiamycin, and oxytetracycline – in varying proportions, along with chlorine dioxide. It is notable, therefore, that out of all four sampled site-years, we find that UIR 2019, which only had hops and chlorine dioxide in its antimicrobial strategy, had a significantly less diverse and more stable microbial composition through time. Even if UIR 2019 did not show increased ethanol output or yield in 2019, relative to 2018 (Supp. Fig. 6), such a stability in the microbial community may be desirable for the industrial process, if it leads to fewer disruptions and/or to a decreased use of antimicrobials, which always add costs and a negative environmental footprint. Limited disclosure of industrial data prevents us from being able to assess other microbiological features that could be informative for this question, such as yeast flocculation and viability.

A large fraction of reads remains unclassified by Kraken 2 (Fig. 2). This is not at all surprising, given the software’s conservative reliance on exact match of k-mers to its database. Therefore, sequence variants (either real or from sequencing error) and low-complexity regions will both lead to unclassified reads. There is a non-negligible chance that the unclassified fraction of reads hides some taxon that is not present in the Kraken 2 database, despite its extensive size. However, given previous knowledge of the microbial composition of the bioethanol fermentation environment, it is unlikely that any such taxon would be more prevalent than *Lactobacillus* or *Limosilactobacillus*.

Our findings underscore the dual-edged role of bacterial contaminants in bioethanol fermentation. We present evidence suggesting that certain *Lactobacillus* species, particularly *L. amylovorus, L. helveticus*, and *L. crispatus*, may exert beneficial effects, potentially enhancing process yield and yeast budding rate at lower abundances. However, exceeding a critical threshold, these same *Lactobacillus* populations are associated with detrimental effects on yeast viability and the occurrence of flocculation, a poorly understood phenomenon in this context that warrants further investigation.

Conversely, our data indicate that *Limosilactobacillus* species (*L. fermentum, L. panis,* and *L. mucosae*) are likely detrimental to fermentation performance. Their association with reduced yeast viability is consistent with their potential to produce inhibitory metabolites such as acetic acid, due to their heterofermentative metabolism.

In conclusion, our metagenomic analysis of industrial sugarcane bioethanol production across two production years and two distinct biorefineries in Brazil reveals nuanced insights into the dynamics and potential impact of microbial contaminants. Contrary to initial expectations, the temporal shifts in the contaminant microbiome within a single harvest appear to be more pronounced than the differences observed between different harvests or between the two mills studied. This highlights the dynamic nature of these microbial communities and the influence of within-season operational variations. A deeper understanding of the complex dynamics between yeast and bacterial communities in industrial bioethanol production holds great potential to optimize process efficiency and reduce the economic losses associated with contamination, opening new ways for novel bioprocess engineering strategies.

Current methods for studying microbial contamination in industrial bioethanol production, such as cultivation-dependent techniques, are limited because they require isolation for identification and cannot accurately quantify diverse bacterial taxa. Similarly, existing cultivation-independent methods often rely on metabarcoding, which only uses sequences of specific DNA regions. Our study introduces a novel approach using whole-metagenome sequencing and analysis, which offers several advantages: it provides access to taxonomic information for a wider range of microorganisms, including viruses and fungi, offers accurate abundance estimation, and achieves high taxonomic resolution down to the genus and species level. This comprehensive strategy allowed us to visualize the distribution of identified microorganisms in UIR and UCP during 2018 and 2019, as shown in Figure 1, specifically focusing on the percentage of *Saccharomyces* species, other fungi, bacteria, and viruses, along with a magnified view of the most abundant bacterial species relative to all identified bacteria.

The spectrum of bacterial genera identified in this study is largely consistent with findings from both historical culture-dependent studies (Gallo, 1898 (13); Rodini, 1985 (15); Neto, 1990 (17)) and more recent cultivation-independent investigations (Bonatelli et al., 2017 (11); Costa et al., 2015 (21); Queiroz et al., 2020 (21); Lino et al., 2021 (23)). Notably, the dominant genera reported in these prior studies, including *Lactobacillus* and *Bacillus*, are also the most prevalent in our metagenomic dataset. While earlier culture-dependent studies identified species such as *Lactobacillus fermentum* (now *Limosilactobacillus fermentum*), *L. helveticus*, *L. plantarum* (now *Lactiplantibacillus plantarum*), *Bacillus subtilis*, *B. coagulans,* and *B. stearothermophilus*, more recent metagenomic studies, including Lino et al. (2021) (23), have provided species-level resolution, identifying *L. amylovorus, L. fermentum, L. helveticus, L. buchneri* (*Lentilactobacillus buchneri*), *L. plantarum*, and *Bacillus cereus* as key players.

Our metagenomic approach allowed for taxonomic classification down to the species level, although the genus level was the primary focus of our analyses, due to the lower confidence and higher proportion of single-species genera. The original Kraken 2 results for read classification beyond genera are included in Supplementary Material 1. Within the *Lactobacillus* genus, *L. amylovorus* was the most abundant species, with *L. helveticus* and *L. crispatus* present at lower levels in some samples. The *Limosilactobacillu*s genus was primarily represented by *L. fermentum*, along with *L. panis* and *L. mucosae*. *Bacillus* was dominated by *Bacillus cereus*, with minor contributions from *Bacillus wiedmannii.* Other identified genera were largely represented by a single classified species: *Lentilactobacillus buchneri, Lactiplantibacillus plantarum, Streptococcus infantarius, Streptomyces sp*. ICC4, *Megasphaera elsdenii, Prevotella sp*. WR041, and *Geobacillus stearothermophilus*. A recent study on bioethanol production identified key bacterial influences on fermentation performance. It found that an increased yeast-to-bacteria ratio correlates with higher production performance. Specifically, *Lactobacillus amylovorus* and *Weissella* species were associated with higher production performance, with *L. amylovorus* potentially having beneficial effects on the ethanol yield. Conversely, *Limosilactobacillus fermentum* was identified as a detrimental bacterium that can reduce ethanol yield by up to 5%. Interestingly, different strains of *L. fermentum* were found to have varying impacts (37).

## CONCLUSION

Intriguingly, the presence of *Bacillus* species (*B. cereus* and *B. wiedmannii*) appears to be negatively correlated with the abundance of both *Lactobacillus* and *Limosilactobacillus*. This suggests a potential for *Bacillus* to act as a natural modulator of the fermentation microbiome, potentially preventing the overgrowth of *Lactobacillus* and the establishment of undesirable *Limosilactobacillus* populations. However, the specific mechanisms underlying this interaction within the complex industrial fermentation environment require further investigation. Interestingly, *B. cereus* was not identified as being part of the taxa that make up 80% of the biodiversity in the samples analysed by Lino et al (2021) (23).

The ability of our shotgun metagenomic approach to provide species-level resolution within key genera like *Lactobacillus* represents a significant advancement over studies that rely solely on genus-level identification. The impact observed even within the *Lactobacillus* genus highlights the importance of future research focusing on strain-specific functional characterization to fully understand their roles in bioethanol production, as had already been shown by Lino et al (2024) (37) and Rich et al (2018) (39). The insights gained from this study pave the way for exploring targeted interventions to manipulate the fermentation microbiome. For instance, further investigation into the antagonistic relationship between *Bacillus* and lactic acid bacteria could lead to the development of novel biocontrol strategies to enhance process stability and efficiency.

To validate the correlations observed in this industrial setting and to understand the underlying mechanisms of interaction between yeast and bacterial contaminants, controlled laboratory experiments and fermentation simulations will be required. Using defined co-cultures with varying proportions of the identified key bacterial species (a.k.a. synthetic microbial communities) would provide a more controlled environment to elucidate the causal relationships between specific microbial populations and fermentation performance, including yield, yeast health, and flocculation dynamics. Furthermore, detailed metabolomic analyses could identify specific microbial metabolites responsible for the observed effects.

Despite the different scopes, this study and that of Rego-Costa et al. (2023) (24) converge on similar ecological patterns. Both show that UCP displays higher microbial diversity, whereas UIR tends to harbor more homogeneous communities, particularly in 2019. Such convergence, emerging from independent analyses of yeast and bacterial populations, suggests that structural factors of the industrial setting (e.g., starter strain choice, recycling practices, or antimicrobial usage) exert a consistent influence on the broader fermentative microbiome. Also, this comparison indicates that, whereas yeast populations undergo rapid and recurrent strain turnover, bacterial contaminants follow more predictable trajectories and appear less affected by yeast dynamics. This distinction may have practical implications: controlling invasive yeast lineages and managing bacterial contaminants may require distinct, yet complementary, strategies to ensure stability and efficiency in industrial ethanol production.

## ACKNOWLEDGMENTS

We thank the various members of the bioethanol mills UCP and UIR who provided extensive support and expertise along the way, especially Thaís Granço and Luciano Zamberlan. We thank the Bauer Core facility at Harvard for assistance with sequencing, as well as the FAS Division of Science Research Computing Group, which manages the computer cluster used for the bioinformatic analysis of the sequencing data.

## SUPPLEMENTARY MATERIAL

This manuscript is accompanied by Supplementary Tables and Figures (3 tables and 11 figures), which can be found in this same document, and by Supplementary Material 1 and Supplementary Material 2, supplied as separate files.

## SUPPLEMENTARY TABLES AND FIGURES

**Supplementary Table 1.** Description of the industrial metrics kindly shared by both UCP and UIR. These metrics primarily serve the internal purposes of these companies, and therefore will have been calculated using undisclosed methods, which may vary from site to site. Therefore, some of the units are unclear or unknown, but the respective metrics were kept in the analysis as they are still interpretable for our purposes. Must is the industry term for the sugary liquid to which yeast is added for fermentation, and wine is the alcoholic broth right at the end of a fermentation cycle.

| Mill | Metric | Description [unit] |
| --- | --- | --- |
| UCP | solids_must | Fraction of sugarcane solids in must [%] |
|  | trs_must | Total reducing sugars in must [%] |
|  | trs_wine | Total reducing sugars in wine [%] |
|  | brix_must | Brix level of must [degrees Brix] |
|  | ktrs_wine | Total reducing sugars in wine, corrected by alcoholic content [unk.] |
|  | kacidity | Acidity of wine, measured by titration [unk.] |
|  | kglycerol | Glycerol content of wine, corrected by unknown factor [%] |
|  | gl_vat | Ethanol content in fermentation vat [degrees Gay-Lussac] |
|  | ph_must | pH of must |
|  | ph_wine | pH of wine |
|  | output | Total weekly ethanol production [liters] |
|  | yield | Net efficiency of production of ethanol from sugar [%] |
| UIR | yeast_viability | Fraction of viable yeasts in fermentation [%] |
|  | budding_rate | Fraction of budding yeast cells in fermentation [%] |
|  | budding_viability | Fraction of viable buds [%] |
|  | floc | Fraction of flocculated cells [%] |
|  | rods | Amount of rod bacteria [rods/ml] |
|  | operation_time | Fraction of time pipeline was operational [%] |
|  | yield | Net efficiency of production of ethanol from sugar [%] |
|  | trs_sugarcane | Total reducing sugars in sugarcane [%] |
|  | output | Total weekly ethanol production [liters] |

**Supplementary Table 2.** Linear regression of UCP industry metrics on *Lactobacillus* and *Limosilactobacillus* abundances. Significance of effect estimates: *: *p* < 0.1, **: *p* < 0.05, ***: *p* < 0.05/12 as a Bonferroni-corrected significance threshold.

| Response | Lactobacillus | | | Limosilactobacillus | | | $R^2$ | Adj. $R^2$ |
| --- | --- | --- | --- | --- | --- | --- | --- | --- |
| | $\beta$ (std. err.) | $t_{df=32}$ | $p$ | $\beta$ (std. err.) | $t_{df=32}$ | $p$ | | |
| solids_must | $5 \times 10^{-3}$ ($1.2 \times 10^{-2}$) | $4.3 \times 10^{-1}$ | 0.672 | $8.9 \times 10^{-3}$ ($1.6 \times 10^{-2}$) | $5.7 \times 10^{-1}$ | 0.573 | 0.05 | -0.03 |
| trs_must | $5.1 \times 10^{-3}$ ($8.2 \times 10^{-3}$) | $6.2 \times 10^{-1}$ | 0.537 | $-8.8 \times 10^{-3}$ ($1.1 \times 10^{-2}$) | $-7.9 \times 10^{-1}$ | 0.435 | 0.06 | -0.02 |
| trs_wine | $3.2 \times 10^{-3}$ ($9.1 \times 10^{-3}$) | $3.5 \times 10^{-1}$ | 0.73 | $-1.5 \times 10^{-2}$ ($1.2 \times 10^{-2}$) | -1.2 | 0.233 | 0.29 | 0.22 |
| brix_must | $8.2 \times 10^{-3}$ ($8.4 \times 10^{-3}$) | $9.7 \times 10^{-1}$ | 0.338 | $-8.1 \times 10^{-3}$ ($1.1 \times 10^{-2}$) | $-7.2 \times 10^{-1}$ | 0.48 | 0.10 | 0.01 |
| kacidity | $4 \times 10^{-2}$ ($9.1 \times 10^{-3}$) | 4.4 | < 0.001*** | $-2.4 \times 10^{-2}$ ($1.2 \times 10^{-2}$) | -1.9 | 0.067* | 0.41 | 0.35 |
| ktrs | $1.6 \times 10^{-3}$ ($8.1 \times 10^{-3}$) | $2 \times 10^{-1}$ | 0.846 | $-1.6 \times 10^{-2}$ ($1.1 \times 10^{-2}$) | -1.5 | 0.144 | 0.21 | 0.14 |
| kglycerol | $-1.1 \times 10^{-2}$ ($1.1 \times 10^{-2}$) | -1 | 0.303 | $1.1 \times 10^{-2}$ ($1.5 \times 10^{-2}$) | $7.4 \times 10^{-1}$ | 0.463 | 0.06 | -0.03 |
| gl_vat | $8.6 \times 10^{-3}$ ($1.1 \times 10^{-2}$) | $7.8 \times 10^{-1}$ | 0.443 | $-8.9 \times 10^{-3}$ ($1.5 \times 10^{-2}$) | $-5.9 \times 10^{-1}$ | 0.557 | 0.22 | 0.15 |
| ph_must | $-9 \times 10^{-3}$ ($1.1 \times 10^{-2}$) | $-8.4 \times 10^{-1}$ | 0.409 | $1.4 \times 10^{-3}$ ($1.5 \times 10^{-2}$) | $9.5 \times 10^{-2}$ | 0.925 | 0.12 | 0.03 |
| ph_wine | $-2.8 \times 10^{-2}$ ($9.1 \times 10^{-3}$) | -3 | 0.005** | $1.5 \times 10^{-2}$ ($1.2 \times 10^{-2}$) | 1.2 | 0.229 | 0.28 | 0.21 |
| output | $-1.3 \times 10^{-2}$ ($9.4 \times 10^{-3}$) | -1.3 | 0.188 | $-6.8 \times 10^{-3}$ ($1.3 \times 10^{-2}$) | $-5.3 \times 10^{-1}$ | 0.599 | 0.07 | -0.02 |
| yield | $3 \times 10^{-3}$ ($2 \times 10^{-3}$) | 1.5 | 0.146 | $2.5 \times 10^{-3}$ ($2.8 \times 10^{-3}$) | $9.1 \times 10^{-1}$ | 0.367 | 0.18 | 0.10 |

**Supplementary Table 3.** Linear regression of UIR industry metrics on *Lactobacillus* and *Limosilactobacillus* abundances. Significance of effect estimates: *: *p* < 0.1, **: *p* < 0.05, ***: *p* < 0.05/9 as a Bonferroni-corrected significance threshold.

| Response | Lactobacillus | | | Limosilactobacillus | | | $R^2$ | Adj. $R^2$ |
| --- | --- | --- | --- | --- | --- | --- | --- | --- |
| | $\beta$ (std. err.) | $t_{df=32}$ | $p$ | $\beta$ (std. err.) | $t_{df=32}$ | $p$ | | |
| yeast_viability | $-2.5 \times 10^{-2}$ ($3.3 \times 10^{-3}$) | -7.3 | < 0.001*** | $-1.4 \times 10^{-2}$ ($6.5 \times 10^{-3}$) | -2.1 | 0.036** | 0.39 | 0.38 |
| budding_rate | $1.5 \times 10^{-2}$ ($4.1 \times 10^{-3}$) | 3.6 | < 0.001*** | $1.1 \times 10^{-2}$ ($7.9 \times 10^{-3}$) | 1.4 | 0.165 | 0.11 | 0.09 |
| budding_viability | $-1.8 \times 10^{-2}$ ($3.7 \times 10^{-3}$) | -4.9 | < 0.001*** | $-3.3 \times 10^{-2}$ ($7.1 \times 10^{-3}$) | -4.6 | < 0.001*** | 0.27 | 0.26 |
| floc | $2.6 \times 10^{-2}$ ($3.6 \times 10^{-3}$) | 7.2 | < 0.001*** | $7.3 \times 10^{-3}$ ($7.1 \times 10^{-3}$) | 1 | 0.3 | 0.29 | 0.27 |
| rods | $2.1 \times 10^{-2}$ ($3.8 \times 10^{-3}$) | 5.5 | < 0.001*** | $2.7 \times 10^{-2}$ ($7.5 \times 10^{-3}$) | 3.6 | < 0.001*** | 0.20 | 0.19 |
| operation_time | $2.4 \times 10^{-3}$ ($4.4 \times 10^{-3}$) | $5.5 \times 10^{-1}$ | 0.587 | $2.6 \times 10^{-2}$ ($8.8 \times 10^{-3}$) | 2.9 | 0.004*** | 0.11 | 0.09 |
| yield | $-8.2 \times 10^{-3}$ ($4.4 \times 10^{-3}$) | -1.9 | 0.066* | $1.5 \times 10^{-2}$ ($8.8 \times 10^{-3}$) | 1.7 | 0.095* | 0.10 | 0.07 |
| atr | $6.8 \times 10^{-3}$ ($4.5 \times 10^{-3}$) | 1.5 | 0.137 | $-8.7 \times 10^{-3}$ ($9.1 \times 10^{-3}$) | $-9.6 \times 10^{-1}$ | 0.34 | 0.05 | 0.02 |
| production | $5.3 \times 10^{-3}$ ($4.2 \times 10^{-3}$) | 1.3 | 0.214 | $3.1 \times 10^{-2}$ ($8.5 \times 10^{-3}$) | 3.7 | < 0.001*** | 0.17 | 0.15 |

**Supplementary Figure 1.**
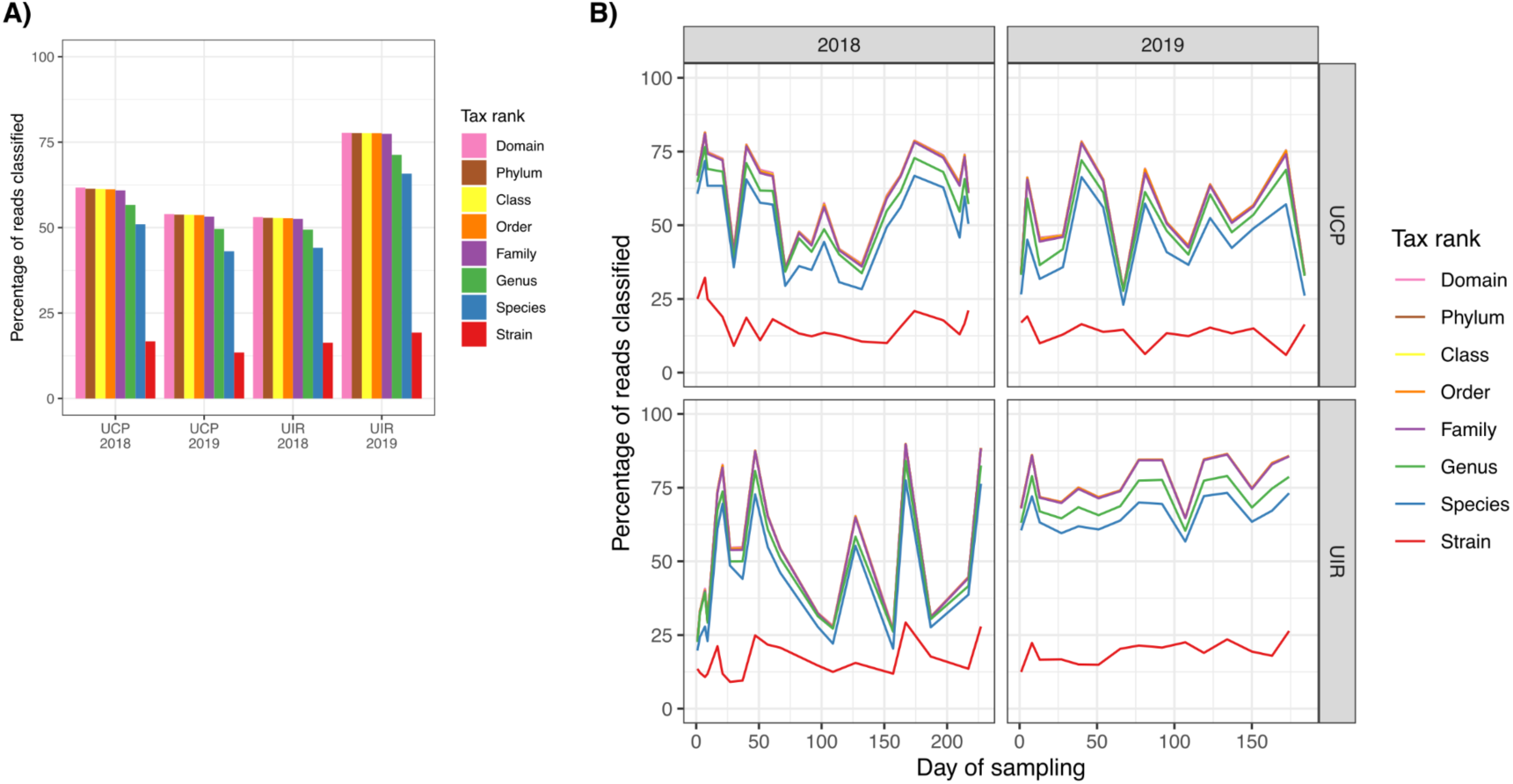
Percentage of reads classified down to different taxonomic levels by Kraken by (A) site-year, and (B) across samples.

**Supplementary Figure 2.**
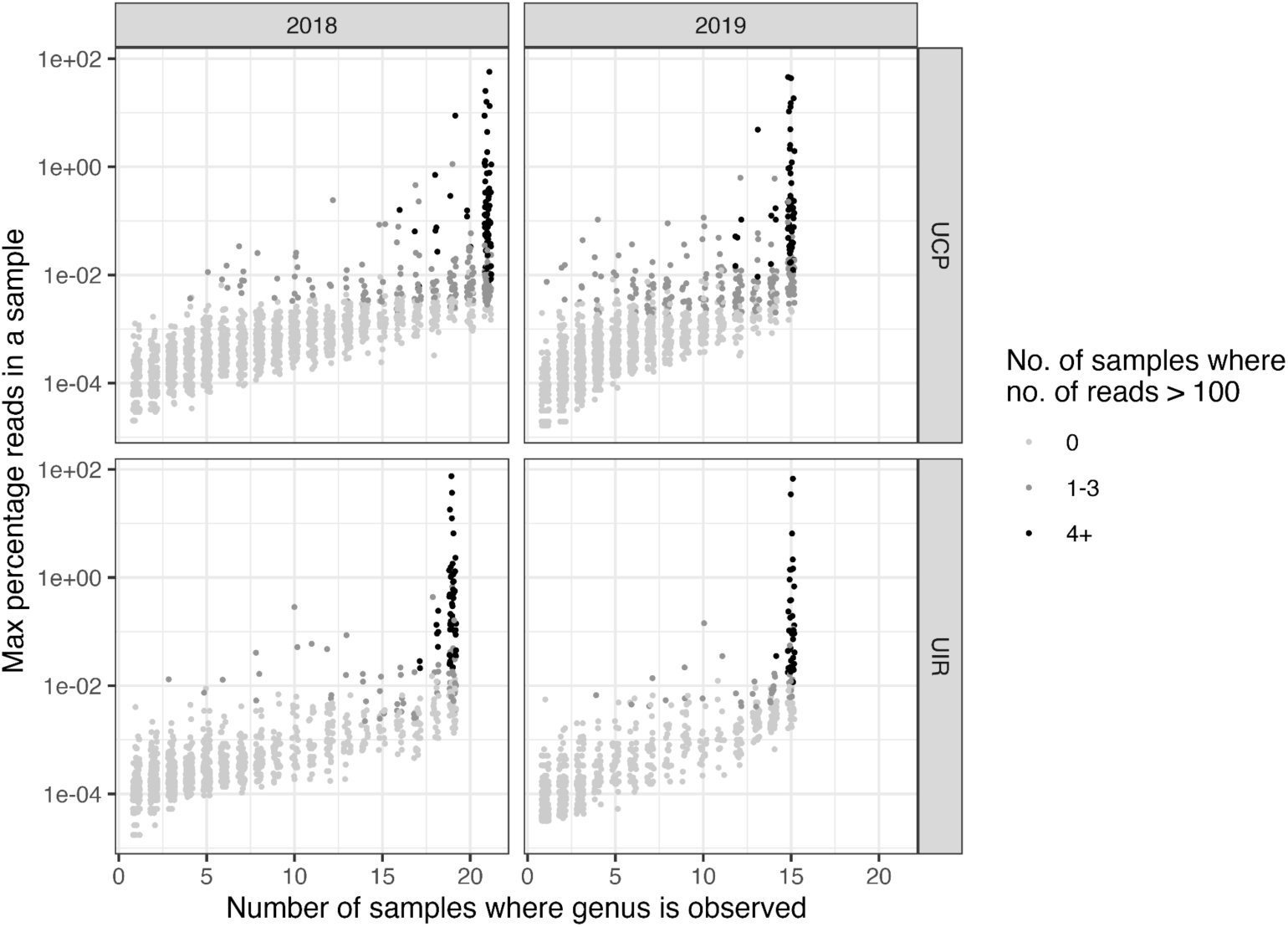
Selection of high-confidence genera across all site-years based on minimum read count and number of samples. Each point represents a genus found at a specific site-year.

**Supplementary Figure 3.**
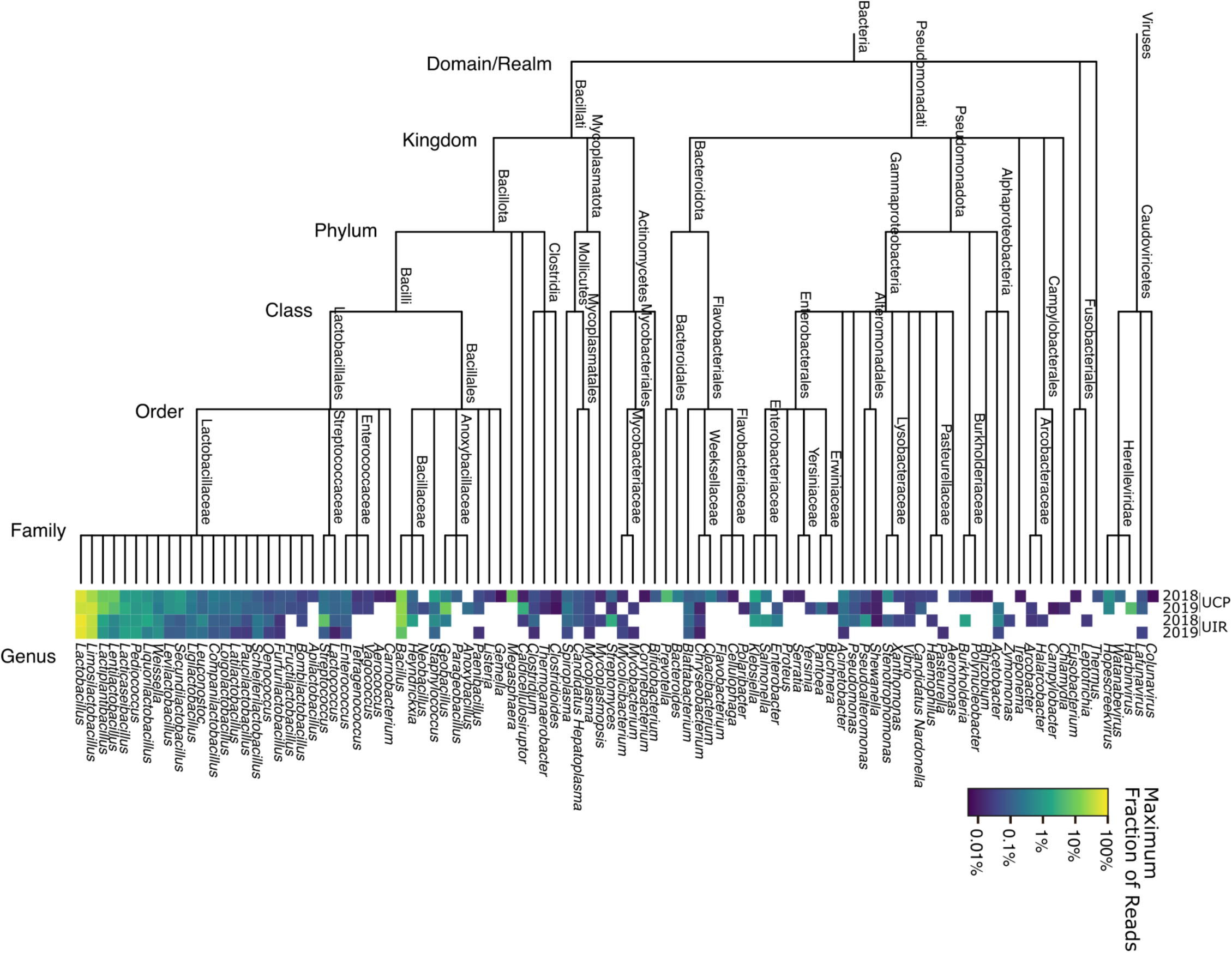
Cladogram of all high-confidence genera selected in the study across all site-years. The maximum abundance of each genus is indicated with a heat map. White tiles in the heatmap indicate genera not observed in that site-year.

**Supplementary Figure 4.**
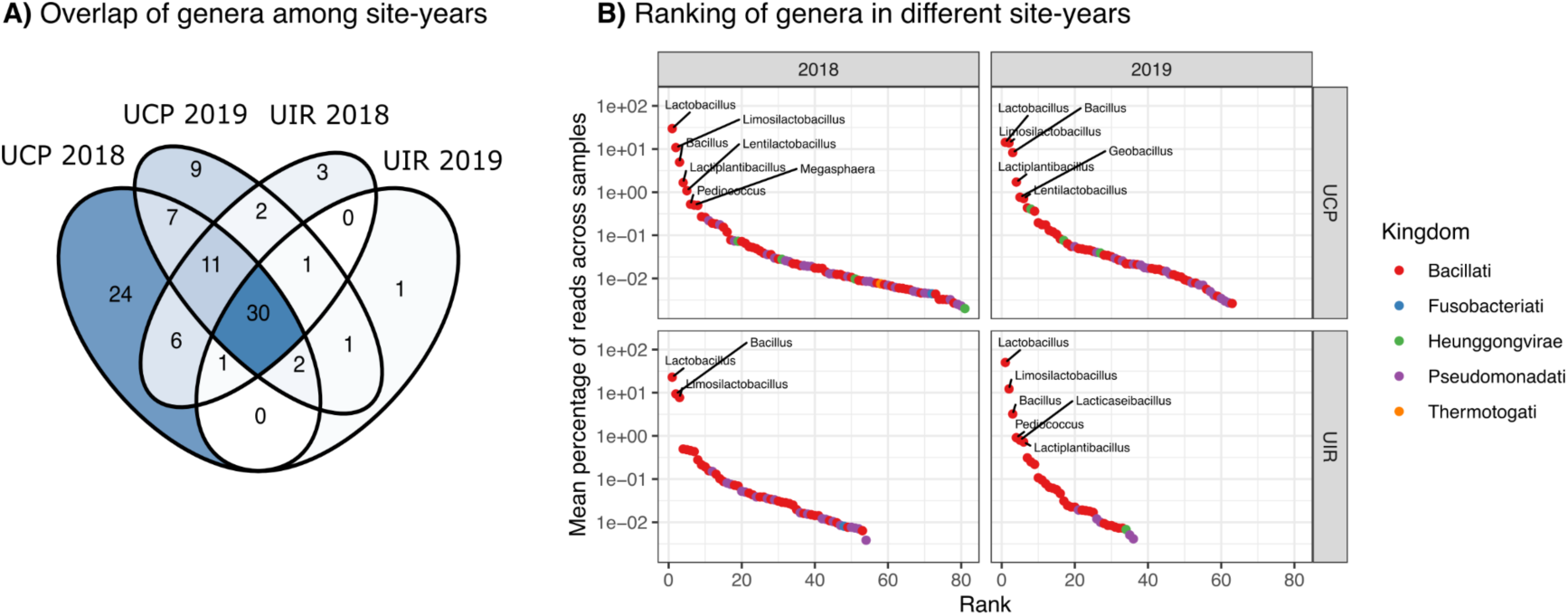
Selected high-confidence genera. (A) Venn diagram of genera among the 4 different site-years. Shading is proportional to the number of genera in each cell. (B) Mean abundance of high-confidence genera selected in each site-year. Each point is a genus, ranked by mean fraction of reads classified to it across all samples in that site-year. Points are colored based on Kingdom classification. Some high abundance genera are labeled.

**Supplementary Figure 5.**
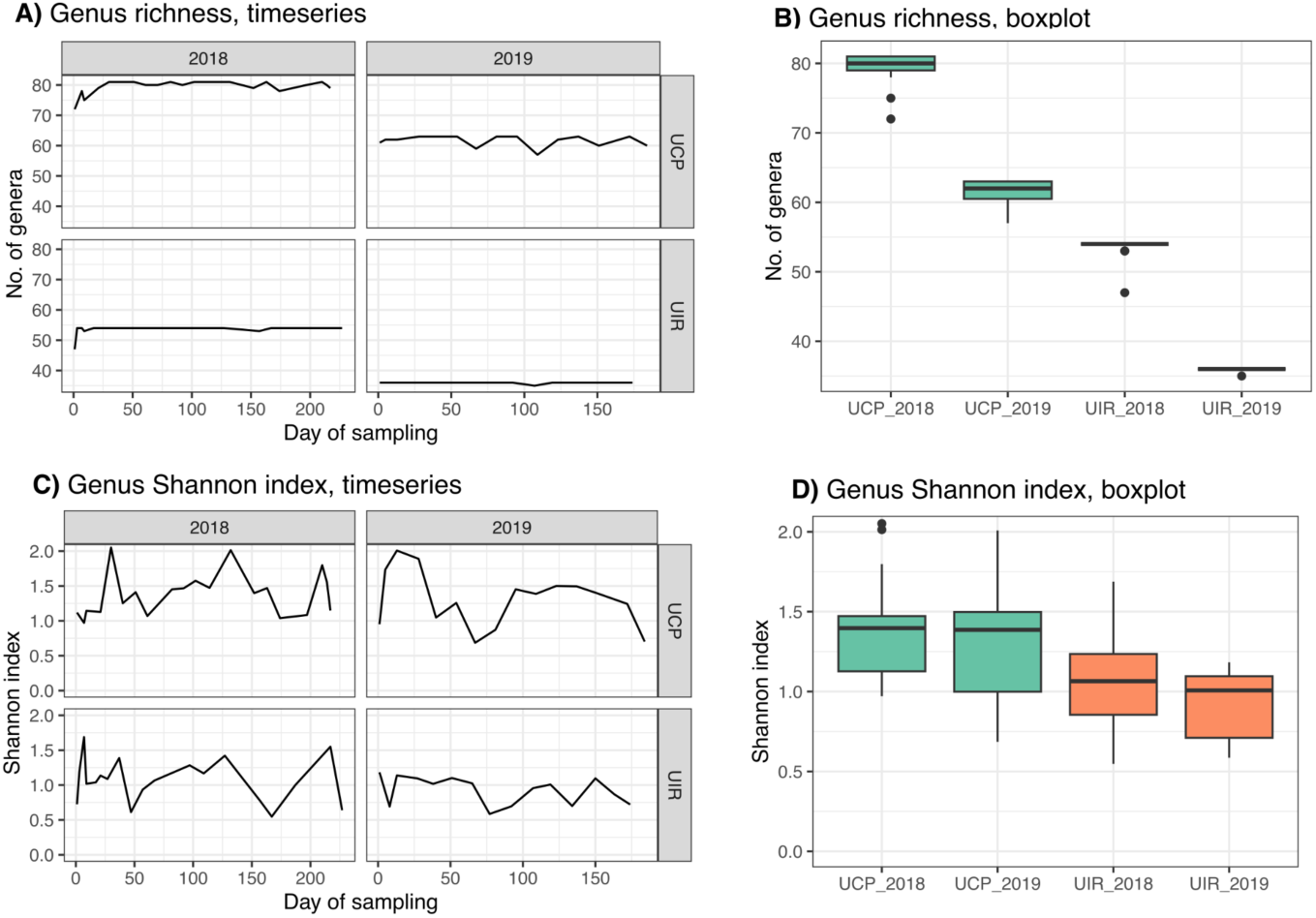
(A, B) Genus richness and (C, D) Shannon index throughout the fermentation season and in each site-year.

**Supplementary Figure 6.**
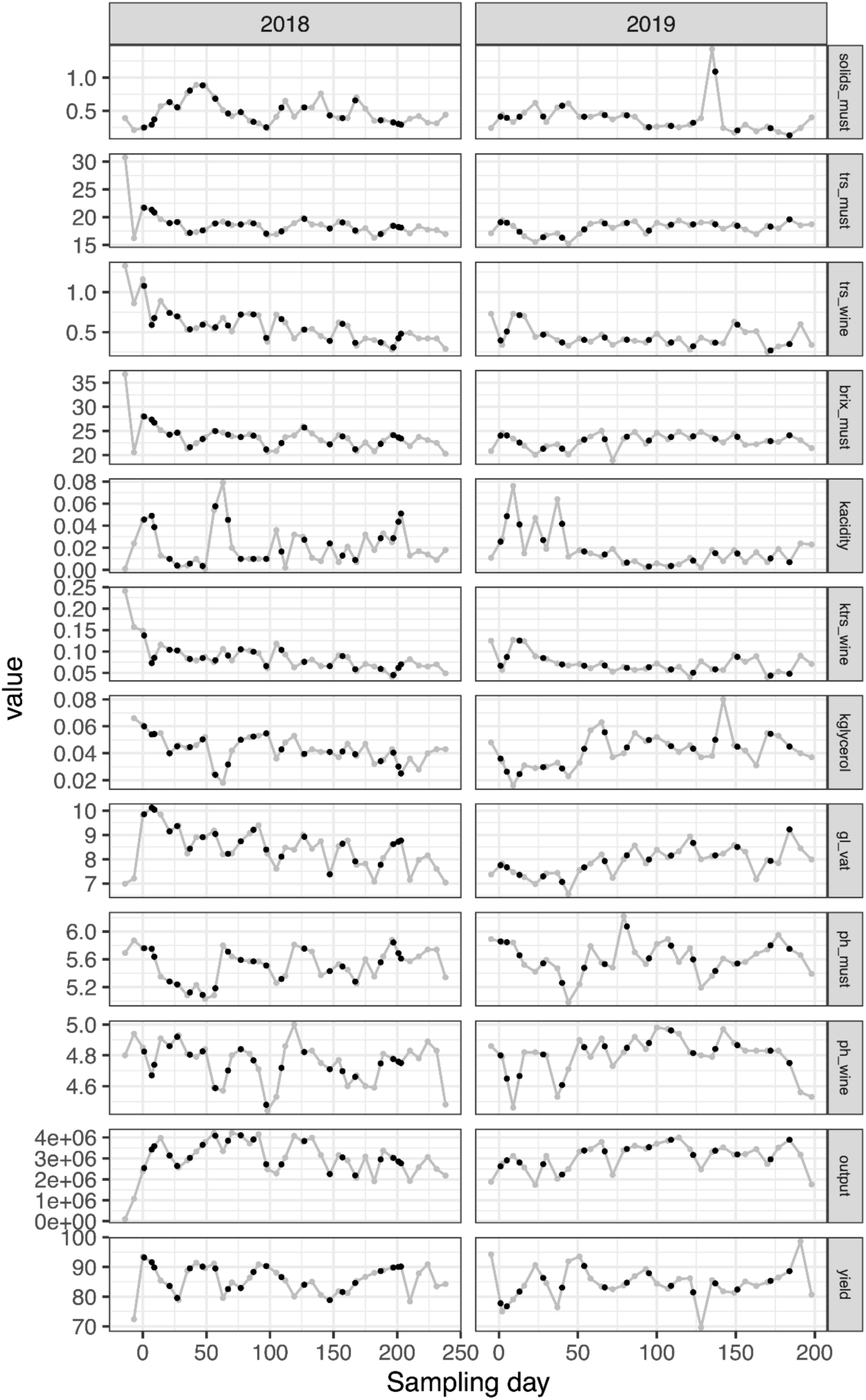
Industrial metrics shared by UCP from the 2018 and 2019 fermentation seasons. Shared data is shown in grey. Linearly-interpolated data points are shown in black and represent estimates of metrics on the days of metagenomic sampling. See Supplementary Table 1 for a definition of the metrics.

**Supplementary Figure 7.**
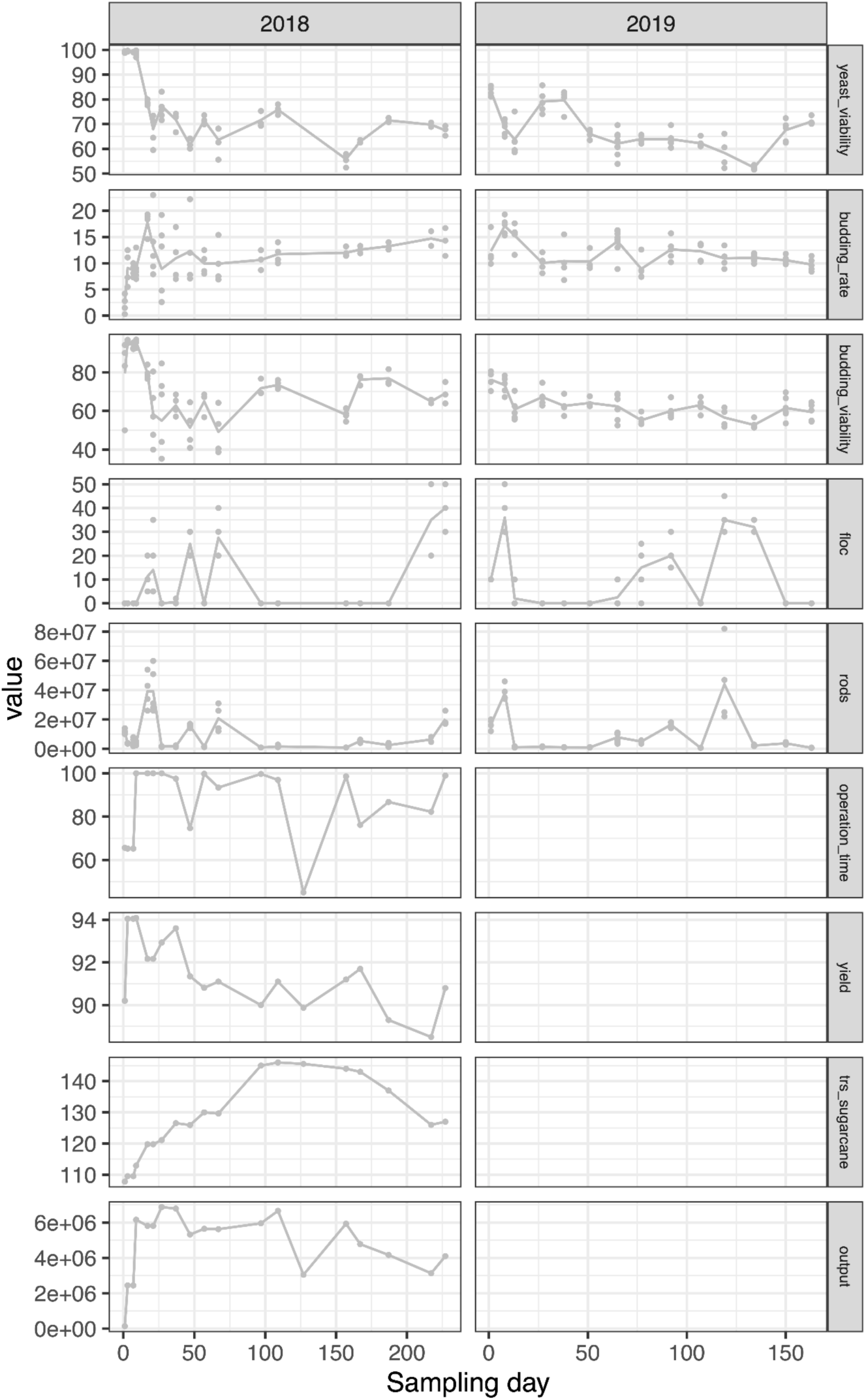
Industrial metrics shared by UIR from the 2018 and 2019 fermentation seasons. Line represents the average value at each day of sampling. See Supplementary Table 1 for a definition of the metrics.

**Supplementary Figure 8.**
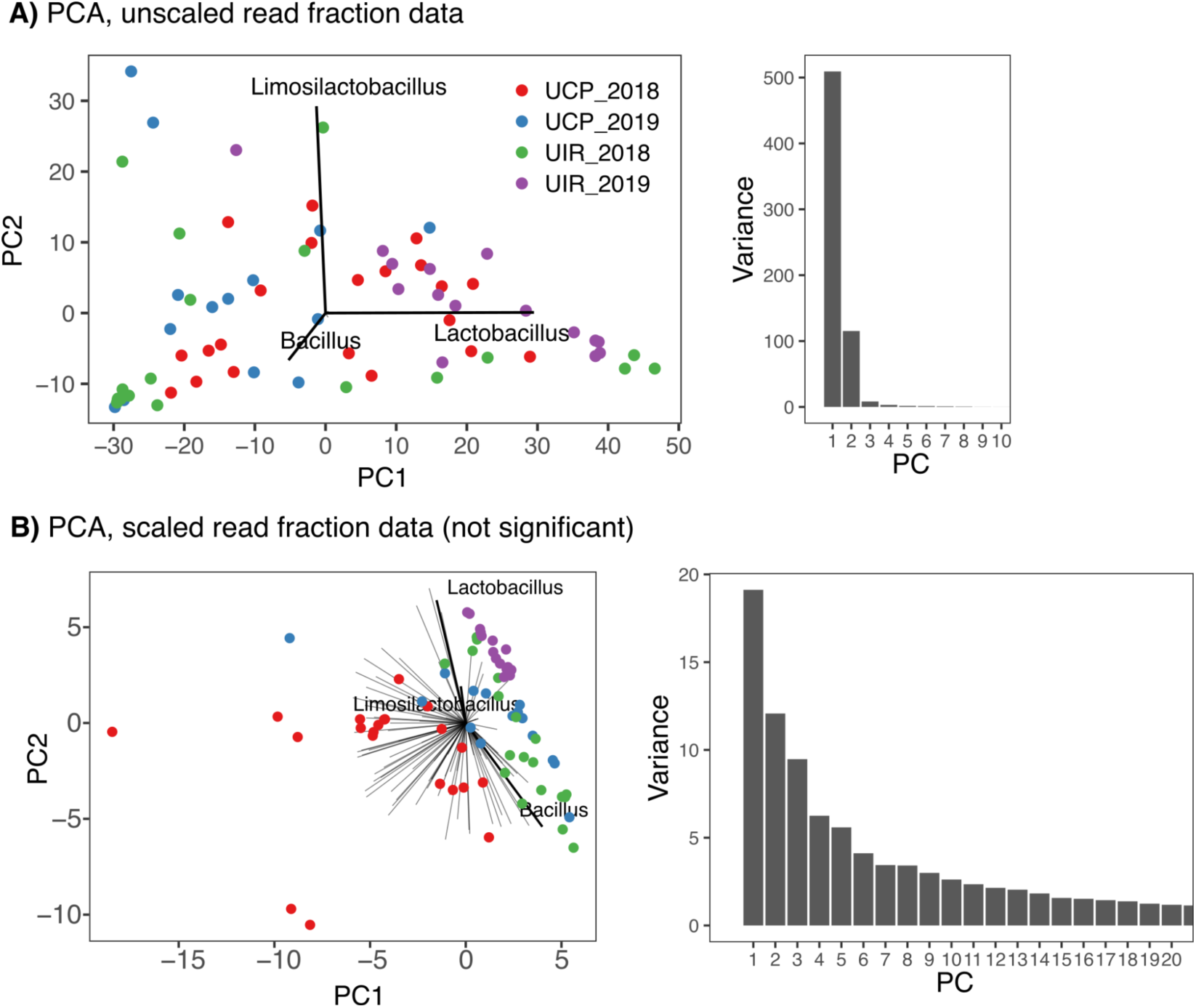
Principal component analysis (PCA) of microbial community composition, using either. (A) unscaled or (B) scaled genus compositional data. Samples are ordinated along the first two principal components (PC1 and PC2), colored by site and year (UCP_2018, UCP_2019, UIR_2018, UIR_2019). The vectors indicate the PCA loadings, interpreted as the taxa whose abundance contributes to each principal component. In the unscaled PCA, *Lactobacillus* and *Limosilactobacillus* each define a PC, after which a few with community variation, including *Bacillus*. The separation of samples along PC1 highlights temporal and site-specific differences in microbial community structure, with UIR_2019 samples clustering distinctly from other groups.

**Supplementary Figure 9.**
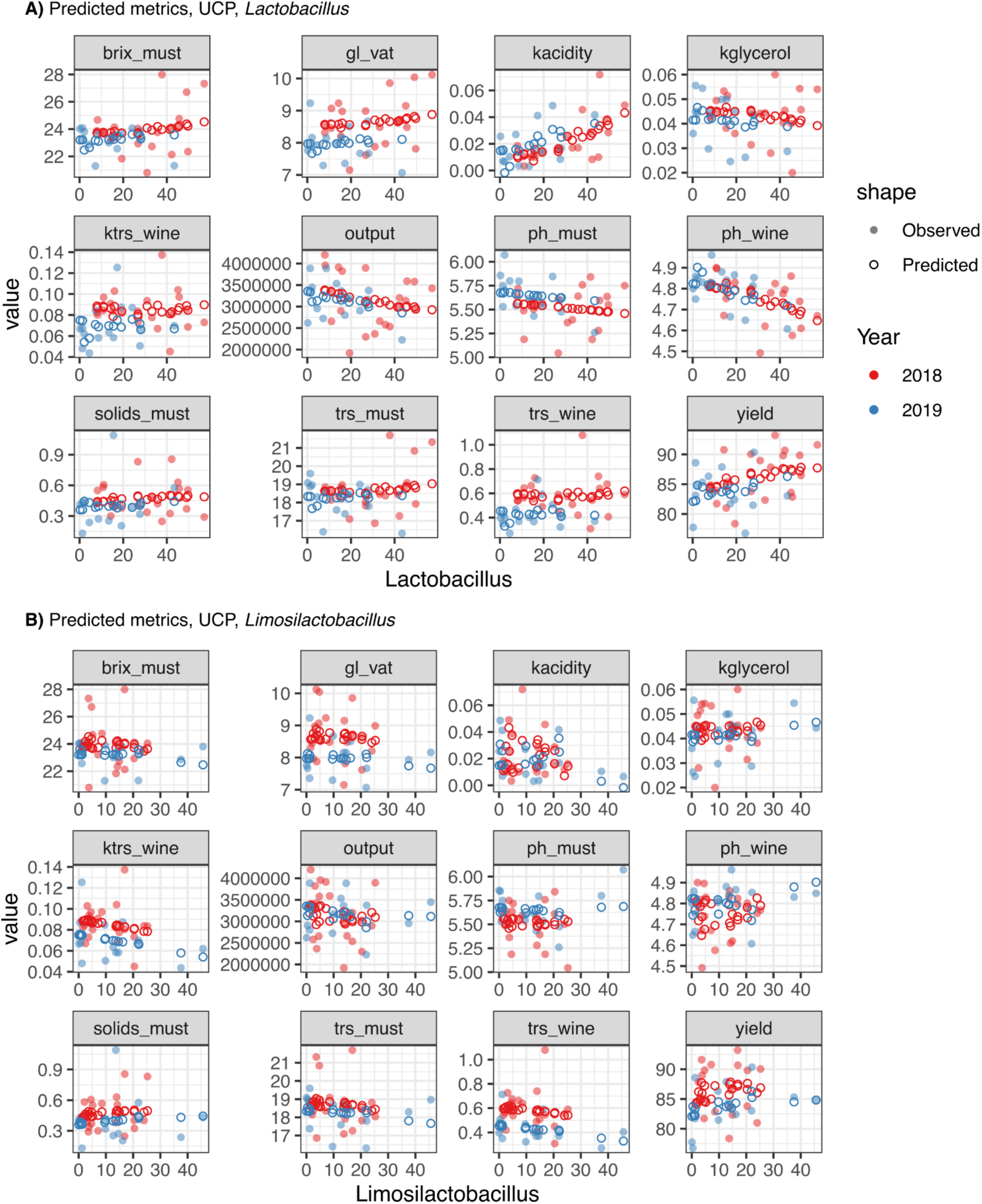
Industrial metrics values against. (A) *Lactobacillus* and (B) *Limosilactobacillus* abundance in UCP. Observed values are shown in light shade full circles, while the corresponding predicted values from the linear regression models are shown in empty circles.

**Supplementary Figure 10.**
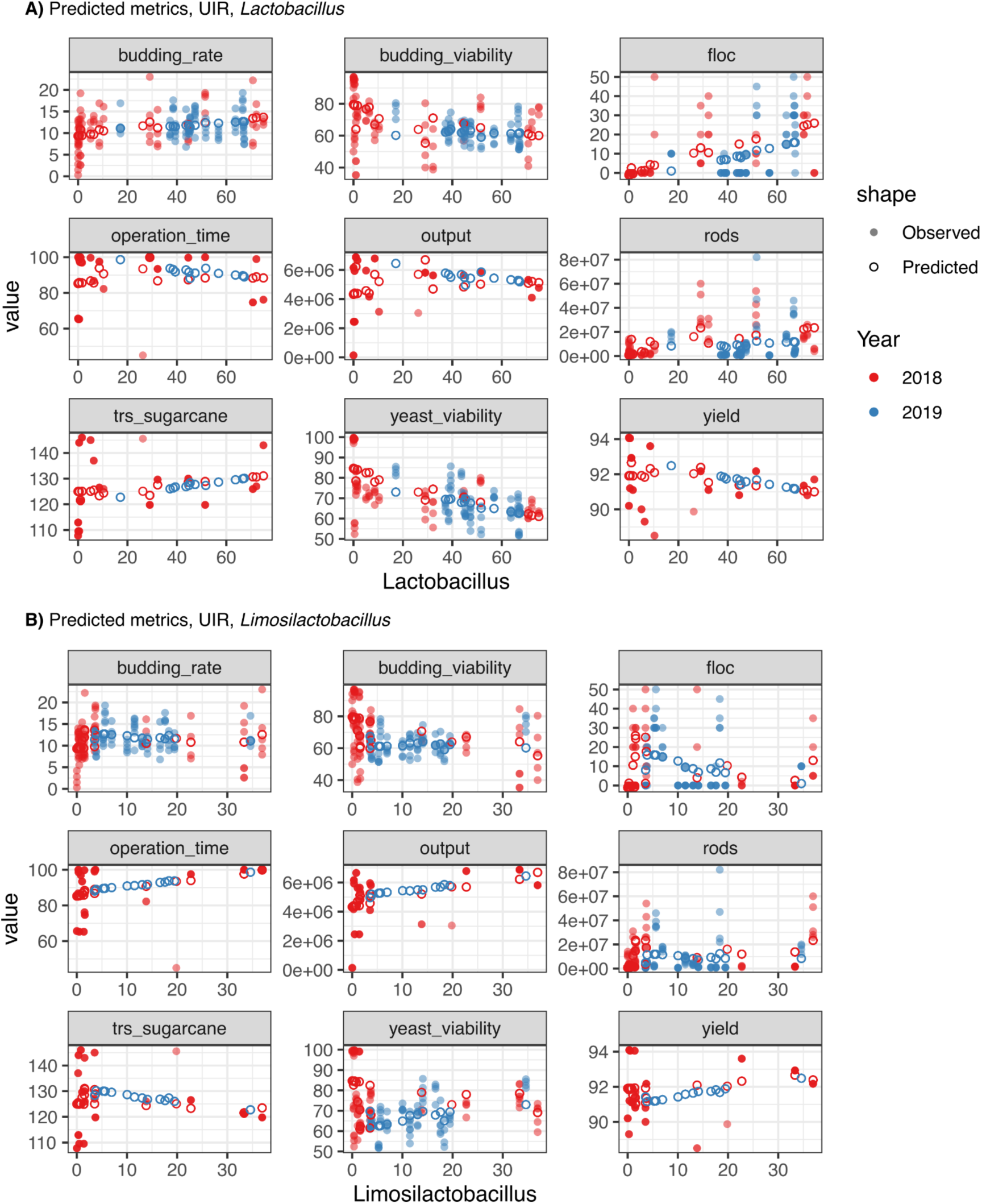
Industrial metrics values against. (A) *Lactobacillus* and (B) *Limosilactobacillus* abundance in UIR. Observed values are shown in light shade full circles, while the corresponding predicted values from the linear regression models are shown in empty circles.

**Supplementary Figure 11.**
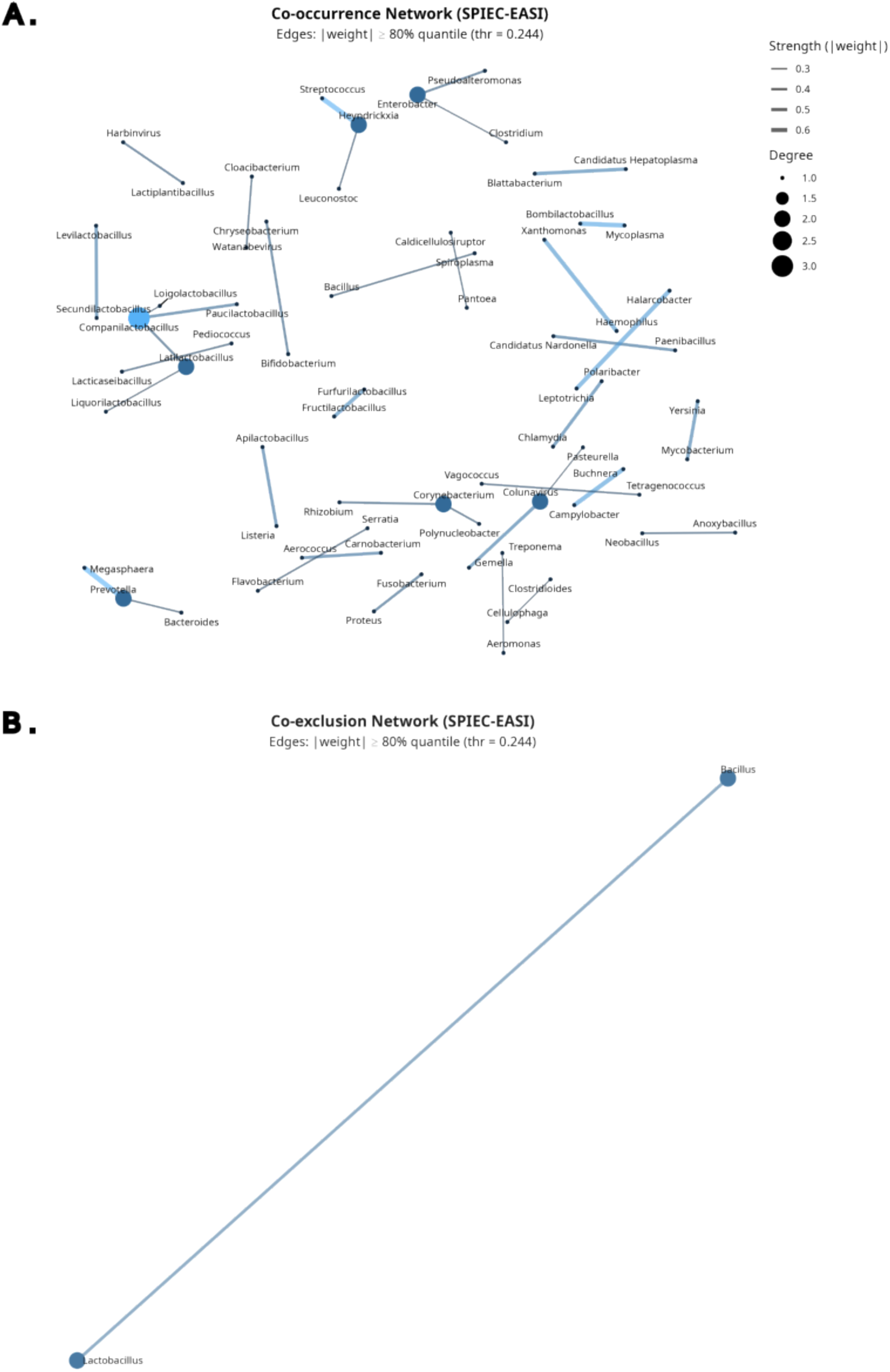
Network analysis of microbial interactions constructed using the SPIEC-EASI. (A) Co-occurrence Network. The topology reveals a decentralized community structure characterized by discrete functional subnetworks rather than a single highly connected core. A prominent subnetwork comprises the newly reclassified *Lactobacillaceae* (e.g., *Companilactobacillus*, *Latilactobacillus*), reflecting shared ecological preferences. Other clusters include soil module (e.g., *Corynebacterium*, *Polynucleobacter*, Rhizobium), and diverse insect/plant commensals. (B) Co-exclusion Network. This subnetwork highlights the *ecological* antagonism between *Bacillus* and *Lactobacillus*.

