## Supplementary Material 2 for "Dynamics of Contaminant Microbes in Bioethanol Production from Sugarcane"

Javascript must be enabled to view this page.

magnitude

UCP\_2018
UCP\_2019
UIR\_2018
UIR\_2019

 100
 100
 99.9999999999999
 100

 52.4452358961862
 42.2787080383021
 44.2347075365872
 69.6684360964132

 52.4452358961862
 42.2787080383021
 44.2347075365872
 69.6684360964132

 .742714404448004
 .291249271716198
 .614948545357288
 .0501825227588759

 .420518305132835
 .208526909745165
 .543014768468905
 .0260094651271275

 .407090531106578
 .168382553680971
 .41584112175938
 .0160459517738663

 .0358452818813137
 .0254505613254352
 .0354785470433531
 .0119321816538125

 .0358452818813137
 .0254505613254352
 .0354785470433531
 .0119321816538125

 .0358452818813137
 .0254505613254352
 .0354785470433531
 .0119321816538125

 .00460933149343253

 .00460933149343253

 .00460933149343253

 .00460502364327552
 .0204237664793018

 .00460502364327552
 .0204237664793018

 .00460502364327552
 .0204237664793018

 .00854423715798259
 0
 .0103444573741113
 .0041137701200538

 .00854423715798259
 0
 .0103444573741113
 .0041137701200538

 .00854423715798259
 0
 .0103444573741113

 0
 0
 0
 .0041137701200538

 .2617442287744
 .0847115886164718
 .258497612296237

 .250303678867415
 .039405907695448
 .258497612296237

 .176287097278348
 .0215392338759362
 .150014692373344

 .0740165815890668
 0
 .0755575849227854

 0
 .0178666738195118
 .0329253350001075

 .00694914098294744

 .00694914098294744

 .00449140892403765
 .0145947289638146

 .00449140892403765

 0
 .0145947289638146

 0
 .0307109519572091

 0
 .00344135217929805

 0
 .0272695997779111

 .0190621697227986
 .00799290772081044
 .0558134629853006

 .012873208757712
 .00498352854436095
 .0519888762886268

 .012873208757712
 .00498352854436095
 .0519888762886268

 .0061889609650866
 .00300937917644949
 .00382458669667376

 .0061889609650866
 .00300937917644949
 .00382458669667376

 .0190752490140299
 .0172588074094911
 .0165601888971015

 .0190752490140299
 .0172588074094911
 .0165601888971015

 .0190752490140299
 .0172588074094911
 .0165601888971015

 .0407369587980015
 0
 .0271038767710796

 .0407369587980015
 0
 .0271038767710796

 .0296518625418127
 0
 .0116963991115744

 .0110850962561888
 0
 .0154074776595052

 .0128680506213437
 .0125449221294608
 .0120429763921976

 .0128680506213437
 .0125449221294608
 .0120429763921976

 .0128680506213437
 .0125449221294608
 .0120429763921976

 .0109056404178219
 0
 .0809620364690854

 .0109056404178219
 0
 .0809620364690854

 .0109056404178219
 0
 .0809620364690854

 .00860173929702585
 0
 .0809620364690854

 .00230390112079606

 .00252213360843483
 .0401443560641941
 .0462116102404389
 .00996351335326121

 .00252213360843483

 .00252213360843483

 .00252213360843483

 0
 .0401443560641941
 .038569197346068
 .00996351335326121

 0
 .0401443560641941
 .038569197346068
 .00996351335326121

 0
 .0401443560641941
 .038569197346068
 .00996351335326121

 0
 0
 .00764241289437095

 0
 0
 .00764241289437095

 0
 0
 .00764241289437095

 .318963426199315
 .0686466207310738
 .0573539621065141
 .0241730576317484

 .238157847783741

 .238157847783741

 .0192017300013417

 .0192017300013417

 .218956117782399

 .218956117782399

 .0808055784155741
 .0686466207310738
 .0573539621065141
 .0241730576317484

 .0808055784155741
 .0686466207310738
 .0573539621065141
 .0241730576317484

 .0257567170229257
 .0545552966748581
 .0500320608990748
 .0191216693581786

 .0257567170229257
 .0545552966748581
 .0500320608990748
 .0191216693581786

 .0130281682813029
 .00709320133968601

 .00505573204597918

 .00797243623532377

 0
 .00709320133968601

 .0420206931113454
 .00699812271652967
 .00732190120743934
 .0050513882735698

 .0198376676249469
 .00699812271652967
 .00732190120743934
 .0050513882735698

 .0221830254863985

 .00323267311585437

 .00323267311585437

 .00323267311585437

 .00323267311585437

 .00323267311585437

 0
 .0102572424207703
 .0145798147818693

 0
 .0102572424207703
 .0145798147818693

 0
 .0102572424207703
 .0145798147818693

 0
 .00788616863723221
 .0145798147818693

 0
 .00788616863723221
 .00700365542638546

 0
 0
 .00757615935548385

 0
 .00237107378353812

 0
 .00237107378353812

 0
 .003818498819189

 0
 .003818498819189

 0
 .003818498819189

 0
 .003818498819189

 0
 .003818498819189

 51.691678993834
 41.9874587665859
 43.6117152954945
 69.6182535736543

 51.5854721047448
 41.8416554870666
 43.3455929335819
 69.566751209685

 51.0977417051382
 41.6426938768897
 43.3142975072903
 69.5583697577645

 45.780747233861
 32.4383567037764
 33.5633886826997
 66.3006384173083

 .00434014719179306

 .00434014719179306

 45.6231277763565
 32.2920940827346
 33.0799636197866
 66.2465928680421

 .00782073548849581

 .00880491010269789
 .0162080930137584
 .014923331212415

 .0779995774246765
 .0550708138054744
 .0290507687594408
 .0462239740826892

 .0142279238718735
 .00700684620855895

 .0124113816327728
 .0122514836173182
 .00684428210162418
 .0093338816443092

 .494517890352746
 .433862322163242
 .456714590450212
 .797263441091397

 1.66506175433812
 1.71148907419494
 .478506580445691
 .722910869237964

 29.7366188659811
 14.2663694640662
 22.9148524485517
 50.1985669442814

 .0490020655584569
 .0425590259491464
 .0195965769903187
 .0224939961254608

 1.08069598961848
 .71075633703681
 .155868234903835
 .249617590147703

 .0720183792159246
 .0460484076352009
 .101898656137893
 .0951485975393648

 .183158557412009
 .174625205783914
 .0383715171972276
 .0646109230274233

 .119352654741362
 .0636796203829579
 .0719217351646443
 .106960833728738

 10.8295040260141
 13.8305718584081
 7.72903625487925
 12.262765182776

 .26075757871661
 .19542704584778
 .191840645546708
 .218739672269117

 .0653249533220489
 .0453824363857656
 .0291424867651535
 .0566428287737558

 .0124466408858216
 .0149393824738264
 .0407643810110089
 .0168736807742379

 .0435640355101162
 .0341112666948098
 .00917876085327145
 .0190511571749708

 .523496866465754
 .360839950683272
 .49617786897635
 .909902751978893

 .0220894419820446
 .0202063432154728
 .0446739512715981
 .0781269613849075

 .154175219601278
 .120239728081772
 .0364257032022497
 .0613230272873918

 .190078328119986
 .130449377086324
 .21417484536601
 .310036554716377

 .00324401458408609

 .00324401458408609

 .0694512216895607
 .0487805809675362
 .0338792699158634
 .0243744414703851

 .0545501495752969
 .0319002794340952
 .0338792699158634
 .0243744414703851

 .00900601844116072
 .0112538103533381

 .00589505367310308
 .00562649118010287

 .08058407403904
 .0974820400742618
 .44954579299726
 .0296711077957985

 .0281686429148259
 .0482489469699139
 .0159632737972239
 .0072927575747711

 .0524154311242141
 .0492330931043479
 .433582519200036
 .0223783502210274

 5.31699447127728
 9.20433717311328
 9.75090882459056
 3.25773134045618

 4.97415885371902
 8.27808471777971
 9.44544829483462
 3.21829367152703

 4.95773458886808
 8.25213963766569
 9.35755328535891
 3.21829367152703

 .0164242648509408
 .0196476547951857
 .0878950094757096

 0
 .00629742531883236

 .00328018001434062

 .00328018001434062

 .0463676639727079
 .785556772176905
 .0270156182795467

 .03567912284347
 .763588297960793
 .0270156182795467

 .0106885411292379

 0
 .0219684742161117

 .00600219580189537

 .00600219580189537

 .0171895206468131
 .0348573094474898
 0
 .00845187874916455

 .0171895206468131
 .0348573094474898
 0
 .00845187874916455

 .269996057122505
 .105838373709178
 .278444911476396
 .0309857901799866

 .269996057122505
 .105838373709178
 .278444911476396
 .0309857901799866

 .00462170523638282
 .176352482483501
 .00633982783511876

 .00462170523638282
 .176352482483501
 .00633982783511876

 .00462170523638282
 .176352482483501
 .00633982783511876

 .00462170523638282
 .176352482483501
 .00633982783511876

 .0294659419322027
 .0226091276933688
 .0249555984564952
 .00838145192056833

 .00328520906098805
 .00246059337742273

 .00328520906098805
 .00246059337742273

 .00328520906098805
 .00246059337742273

 .0199693768808285
 .0170592668722506
 .0249555984564952
 .00838145192056833

 .0199693768808285
 .0170592668722506
 .0249555984564952
 .00838145192056833

 .0199693768808285
 .0170592668722506
 .0249555984564952
 .00838145192056833

 .00621135599038619
 .00308926744369543

 .00621135599038619
 .00308926744369543

 .00621135599038619
 .00308926744369543

 .453642752437942

 .453642752437942

 .453642752437942

 .453642752437942

 .0366260844750496
 .0202218714724916
 .143488631185877
 .0256755258042072

 .0366260844750496
 .0202218714724916
 .143488631185877
 .0256755258042072

 .0056559282603073

 .0056559282603073

 .0056559282603073

 .0093162982585035
 .0108565955704665
 .014162062106509
 .0256755258042072

 .0026953662685453

 .0026953662685453

 .0066209319899582
 .0108565955704665
 .014162062106509
 .0256755258042072

 .0066209319899582
 0
 .014162062106509
 .0179735677384637

 0
 .0108565955704665
 0
 .0077019580657435

 .0216538579562388
 .00936527590202507
 .129326569079368

 .0216538579562388
 .00936527590202507
 .129326569079368

 .0216538579562388
 .00936527590202507
 .129326569079368

 .0695808046141397
 .125581408046877
 .122633730726774
 .0258268381650839

 .0641008715070697
 .125581408046877
 .111761546577732
 .0258268381650839

 .0255511307299645
 .0429672107420813
 .042656241854627
 .00729316898228081

 .0149616635961038
 .0266903958848525
 .0284077767549005

 .0149616635961038
 .0266903958848525
 .0284077767549005

 .0105894671338607
 .0162768148572288
 .0142484650997265
 .00729316898228081

 .0105894671338607
 .0162768148572288
 .0142484650997265
 .00729316898228081

 .0385497407771052
 .0826141973047957
 .0691053047231055
 .0185336691828031

 .0385497407771052
 .0826141973047957
 .0691053047231055
 .0185336691828031

 .0385497407771052
 .0826141973047957
 .0691053047231055
 .0185336691828031

 .00547993310707003
 0
 .0108721841490413

 .00547993310707003
 0
 .0108721841490413

 .00547993310707003
 0
 .0108721841490413

 .00547993310707003
 0
 .0108721841490413

 .00441940591780887
 0
 .00804369573533124

 .00441940591780887
 0
 .00804369573533124

 .00441940591780887
 0
 .00804369573533124

 .00441940591780887
 0
 .00804369573533124

 .00441940591780887

 .00441940591780887

 0
 0
 .00804369573533124

 0
 0
 .00804369573533124

 .00642309198642228

 .00642309198642228

 .00642309198642228

 .00642309198642228

 .00642309198642228

 .00642309198642228

 .0931241460682197
 .456087188052157
 0
 .00632082186177198

 .0931241460682197
 .456087188052157
 0
 .00632082186177198

 .0931241460682197
 .456087188052157
 0
 .00632082186177198

 .0931241460682197
 .456087188052157
 0
 .00632082186177198

 .0931241460682197
 .456087188052157
 0
 .00632082186177198

 .00162077366757218

 .00162077366757218

 .00162077366757218

 .0836057255815297
 .424929454422137

 .0836057255815297
 .424929454422137

 .0624741488596352
 .0757156465712354

 .0211315767218945

 0
 .349213807850902

 .0078976468191178
 .0311577336300194
 0
 .00632082186177198

 .0078976468191178
 .0311577336300194
 0
 .00632082186177198

 .0078976468191178
 .0311577336300194
 0
 .00632082186177198

 47.4616399577456
 57.2652047736457
 55.7652924634128
 30.325243081725

 47.4616399577456
 57.2652047736457
 55.7652924634128
 30.325243081725

 47.4616399577456
 57.2652047736457
 55.7652924634128
 30.325243081725

 47.4616399577456
 57.2652047736457
 55.7652924634128
 30.325243081725

 47.4616399577456
 57.2652047736457
 55.7652924634128
 30.325243081725

 47.4616399577456
 57.2652047736457
 55.7652924634128
 30.325243081725

 47.4616399577456
 57.2652047736457
 55.7652924634128
 30.325243081725

 47.4616399577456
 57.2652047736457
 55.7652924634128
 30.325243081725
